# Bumblebee string-pulling skill spreads between colonies under open diffusion conditions

**DOI:** 10.1101/2025.03.03.641175

**Authors:** Olga Procenko, Alice D. Bridges, Amelia Kowalewska, Mikko Juusola, Lars Chittka

**Affiliations:** School of Biological and Behavioural Sciences, Queen Mary University of London, United Kingdom; Current address: School of Biosciences, University of Birmingham, Birmingham, UK; Current address: School of Biosciences, University of Sheffield, Sheffield, UK; Current address: Neuroscience Institute, University of Sheffield, Sheffield, UK

## Abstract

Socially-transmitted behavioural traits can, if they persist in a group of animals over time, give rise to locally-adapted phenotypes that can enhance survival. This capacity is widespread through the animal kingdom, and forms the foundation of cultural inheritance. While social learning is well-documented among insects, and particularly in social insects such as bumblebees, the extent to which such behaviours can spread beyond initial kin groups and persist over time remains largely unknown. String-pulling is a non-natural foraging behaviour where bees must manipulate a string to extract an out-of-reach artificial flower and collect a reward, and has previously shown to spread via social learning. However, this was demonstrated only in highly controlled paired-dyad settings, where interactions between bees were strictly limited. Here, we show that string-pulling can spread both within and between bumblebee colonies and persist over time, under previously-untested open diffusion conditions. These are of greater ecological validity compared with classical paired dyad paradigms, and involve the seeding of a manually-trained demonstrator into a group of naïve conspecifics. From this single point of origin, string-pulling behaviour spread rapidly within original, ‘primary’ colonies. Once the behaviour was established in the primary colonies, ‘secondary’ colonies were introduced, and string-pulling was also acquired by these new foragers. Furthermore, string-pulling was acquired through individual trial-and-error learning by a small number of bees in control colonies, which lacked trained demonstrators. These results confirm and build upon previous findings in bumblebees, and contribute to a growing body of evidence suggesting that social learning enables animals to establish local behavioural adaptations in the absence of the computational power provided by large brains.

## Introduction

In order to survive in any particular habitat, organisms must adapt to suit the specific demands and constraints of that habitat: from its climate to the availability of its resources, and to other species sharing it. This generally results in alterations to an organism’s phenotype: its physical appearance, developmental processes, or behaviour. While some traits are genetically predetermined and shaped over multiple generations, and thus may not keep pace with increasingly rapid environmental change, others are flexible and influenced by the environment. One such adaptive trait is behaviour, where behavioural plasticity provides a faster, more flexible ways for animals to adapt (Snell-Rood, 2013). In particular, altering one’s behaviour can allow the exploitation of novel resources that might appear in one’s environment. In the United Kingdom, blue tits learned to peck through foil milk bottle-tops to steal the cream (Fisher and Hinde, 1949). In Australia, cockatoos discovered how to break into rubbish bins and overcame many human attempts to stop them (Klump *et al*., 2021, 2022).

At least in the first instance, novel behaviours such as these had to emerge through individual trial-and-error learning. However, individual learning consumes time and energy, and can be risky. In contexts such as foraging on novel items or interacting with new species, making the wrong decision can be fatal (Laland, 2004). However, when the end result of this trial-and-error is acquired by the innovator’s conspecifics via social learning, it can result in the rapid spread of beneficial new behaviour throughout a population. When this persists over time, it essentially acts as a behavioural “tradition” (Fragaszy and Perry, 2003) and can be considered a unit of *culture.* Often termed a ‘second form of inheritance’ (Whiten, 2017), and once thought wholly unique to humans, culture-like phenomena are increasingly understood to be prevalent throughout the animal kingdom. Chimpanzees from one location use stone tools to crack nuts while their neighbours don’t, for example (Boesch *et al*., 1994; Gunasekaram *et al*., 2024); while male humpback whales change their songs to follow new trends (Garland *et al*., 2011). The majority of studies concerning social learning and culture in animals have focused on vertebrates with relatively large brains, as these plus the ‘complex’ cognitive abilities they support were deemed vital prerequisites (Boyd and Richerson, 1996). However, similar phenomena have since been demonstrated in a very different species: the bumblebee, *Bombus terrestris* (Bridges *et al*., 2023). If opportunity arises, bumblebees too are capable of sustaining their own local variations in their behaviour: in this case, opening an artificial flower in the form of a puzzle-box in one of two possible ways, with dynamics mirroring those previously observed in chimpanzees, vervet monkeys and great tits (Whiten, Horner and de Waal, 2005; van de Waal, Borgeaud and Whiten, 2013; Aplin *et al*., 2015).

*B. terrestris* is a primitively eusocial insect that lives in small colonies of ∼160 individuals (Goulson *et al*., 2002), and is an excellent model for social learning studies in the laboratory. These bumblebees readily learn novel, non-natural foraging behaviours such as pulling strings in exchange for rewards, either through incremental training by an experimenter or through social learning from their trained conspecifics (Alem *et al*., 2016). However, the spread of this behaviour was not tested under so-called “open diffusion” conditions (Whiten and Mesoudi, 2008). Open diffusion paradigms involve the seeding of a trained individual into a larger group of animals, who are then all permitted to interact freely with the substrates necessary to perform the target behaviour with minimal interference from the experimenter. They are considered to be of greater ecological validity when compared with paired dyad paradigms, and better represent the spread of novel behaviour in a group of animals as a local adaptation. Thus, in the present study, we sought to build on our previous work on bumblebee social learning by monitoring the spread of string-pulling under open diffusion conditions. In particular, we investigated whether the spread of string-pulling would be limited to a single colony, or could spread beyond these limits to new, naïve colonies after being established in just one. Demonstrating between-colony transmission would confirm that cultural traits can move beyond the confines of a single, genetically-related group in bumblebees, suggesting that these behaviours are unlikely to represent isolated phenomena and can spread beyond localised groups. Furthermore, our finding that some bumblebees were able to acquire string-pulling in the absence of a trained demonstrator mirrored previous results with our puzzle-box paradigm (Bridges *et al*., 2023), and confirmed the impressive capacity for behavioural flexibility in bumblebees. Taken together, our results highlight and characterise a possible route through which novel foraging adaptations could naturally emerge and become established among bumblebee populations in the wild.

## Materials and methods

### Animal model

Commercially-raised *Bombus terrestris* (spp. audax) colonies (n=9) were supplied by Agralan, Ltd. (Swindon, UK). Upon arrival, each colony was transferred to bipartite wooden nest boxes (40.0 × 28.0 × 11.0 cm), consisting of a covered nest chamber and a vestibule. During transfer, all individuals were marked with unique numbered Opalith tags, to enable accurate identification and tracking of individual behaviour throughout the experiment. This was achieved by trapping each bee in a specially-designed cage, lightly pressing it against a mesh ceiling with a sponge, and fixing the tag to the thorax with superglue. Bees that eclosed during the experiments were also tagged once they attempted to leave the nest. During the experiment, bees were fed with commercially-available pollen every two days, and sucrose solution (20% w/w) every day, and were maintained at standardised room temperature.

Colonies C1, C2 and C3 were seeded with trained demonstrators and used as primary experimental colonies during the first phase of the experiment (*Phase I*), while colonies CC1, CC2 and CC3 served as controls. Colonies SC1, SC2, and SC3 were used as secondary colonies and introduced to the primary colonies during *Phase II* of the experiment.

### Ethical note

All experiments were conducted by researchers trained to handle and maintain bumblebees, and every effort was taken to minimise stress to the subjects. Bumblebees were never made to take part in experiments on the same day that they had been marked: although this process was non-invasive, they were allowed to recover and acclimate in their nest box without further interference. In cases where bees were tagged during the course of an ongoing open diffusion experiment, they were permitted to leave the nest and participate if they so chose but were not ‘encouraged’ to do so. Nest box vestibules were regularly cleaned of debris and waste, and colonies had free access to food outside the daily experimental period. As these were commercially-bred bumblebee colonies, it was not possible for them to be released to the wild after use in our experiments: thus, colonies were euthanised by freezing when they showed signs of aging and collapse. We also aimed to use the minimum number of colonies possible for this experiment.

### Experimental set-up and artificial flower design

Each nest box was connected to the experimental flight arena (120.0 × 100.0 × 37.0 cm; **Fig. 1A**) via a transparent Plexiglas corridor (25.0 × 3.5 × 3.5 cm), which had plastic sliding stoppers inserted into 1.0 mm-wide gaps along its length. These controlled which bees were able to enter the flight arena. The flight arena itself was covered with a transparent Plexiglas lid that prevented bees from escaping and permitted close observation, and was bipartite: a removable central cardboard wall created two separate experimental arenas. This allowed simultaneous experiments to be conducted.

**Figure 1.**
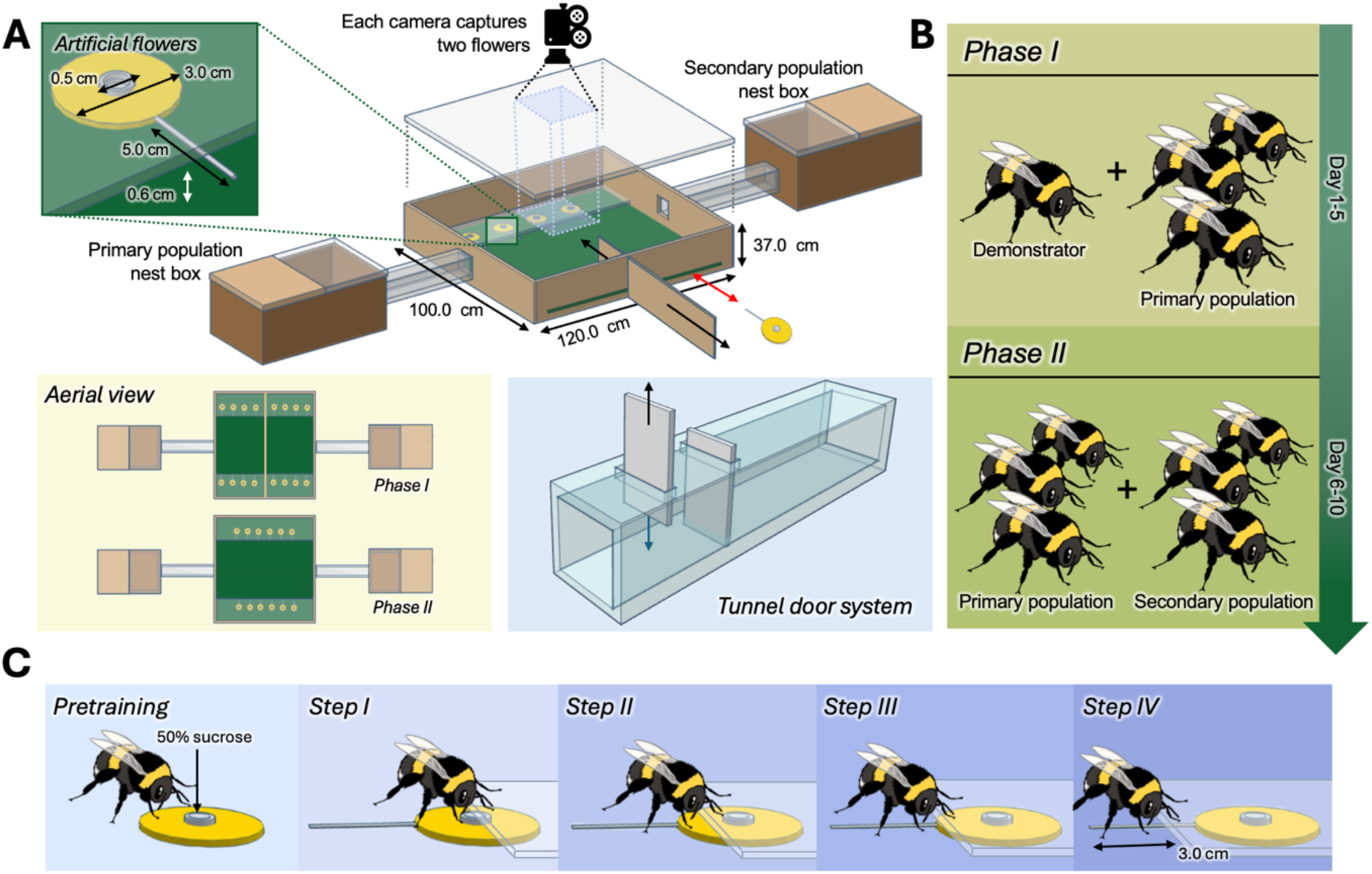
Experimental set-up and protocols. **(A) Experimental set-up.** The experimental arena was made bipartite by a removable divider, allowing for the connection of multiple colonies and the running of simultaneous experiments. Nest boxes were attached to the arena via acrylic tunnels, and a sliding door system (see inset) was used to control access to the arena. Either n=8 or 12 artificial flowers were presented to the bees during *Phase I* and *II* experiments, respectively, and could be removed from the arena by the experimenters through a small gap cut into the sides of the box. Cameras were positioned on top of the clear arena lid, and each field of view captured two of the artificial flowers. **(B) Schematic of experimental phases.** Phase I involved the training of a demonstrator by experimenters, and then tracking the spread of this behaviour throughout the demonstrator’s home colony (the ‘primary population’) under open diffusion conditions for 5 days. Control colonies were exposed to the artificial flowers for the same length of time, but no trained demonstrator was added. Phase II involved the introduction of another colony (the ‘secondary population’), which was allowed to forage alongside the primary colony for a further 5 days. **(C) Stepwise demonstrator training protocol, as described in Alem *et al.,* 2016.** All bees used in the experimental and control open diffusion experiments underwent the pretraining phase, to familiarise them with foraging on the yellow artificial flowers. Once an individual had been selected for demonstrator training, it proceeded through steps I-IV in isolation from other bees. In step I, 50% of the flower was positioned beneath the Plexiglas sheet, with the sucrose reservoir partially accessible. In step II, 75% of the flower was positioned beneath the Plexiglas, forcing bees to physically move the flower to obtain the reward. In step III, the flower was positioned entirely beneath the Plexiglas, and in step IV, it was moved even further back, with ∼3.0cm of the string being the only protruding element.

The artificial flowers (**Fig. 1A inset**) consisted of yellow polyurethane discs (Ø 3.0 cm), with a central sucrose reservoir made from an Eppendorf cap (Ø 0.5 cm). A 5.0 cm-length of cotton string was hot-glued to the edge of each flower. These flowers were placed under a transparent sheet of Plexiglas (6.0 cm wide x 120.0 cm long), so that they could be observed at all times by the experimental bees. This Plexiglas sheet was positioned 0.6 cm above the arena floor to prevent bees squeezing beneath it and reaching the reward without string-pulling. A 2.0 cm-tall hole was cut into the side of the box to allow the flowers to be removed, refilled and replaced from the outside, without needing to remove the flight arena lid, and the Plexiglas sheet over the flowers was angled upwards so that it covered this gap.

### Experimental protocol

Following initial acclimatisation to the arena and artificial flowers via group pretraining, and the selection and training of a demonstrator, the cultural transmission experiment was divided into two main phases as follows. *Phase I* involved the introduction of a trained demonstrator into a colony of naïve foragers (the “primary population”), allowing us to observe the spread of the string-pulling behaviour through social learning and compare it to control colonies with no demonstrator. In *Phase II*, a secondary colony of naïve bees (the “secondary population”) was introduced to forage alongside the primary population, testing the transmission of learned behaviour across colony boundaries (**Fig. 1B**). All experiments were conducted under standardised artificial lights (12:12, high-frequency fluorescent lighting; TMS 24F lamps with HF-B 236 TLD [4.3 kHz] ballasts [Koninklijke Phillips NV, Amsterdam, the Netherlands], fitted with Activa daylight fluorescent tubes [OSRAM Licht AG, Munich, Germany]).

*Initial acclimatisation and group pretraining*. This process was conducted for each colony, to ensure all bees were familiarised with the arena and needing to forage there. First, for 24 h, all bees were permitted to freely enter and forage from a central gravity feeder (containing 20% w/w sucrose solution). Following this, eight artificial flowers without strings attached were randomly positioned throughout the arena; each containing 50 μl 50% w/w sucrose solution. Bees were permitted to forage on these flowers for a further 8 h. This not only allowed bees to associate the yellow flowers with reward, but also the identification of suitable candidates for demonstrator training. Only those individuals that made repeated, consecutive foraging trips throughout this 8 h period were selected to undergo demonstrator training.

*Stepwise demonstrator training protocol*. One individual per *Phase I* experimental colony was selected to undergo demonstrator training, with all other bees restricted to the nest box to prevent familiarisation with the task. Training was performed using the protocol previously described for the original dyadic transmission chain experiments (Alem *et al*., 2016). In brief, the selected bee was presented with artificial flowers (n=4), each attached to a string with 50 μl 50% w/w sucrose in the reservoir. However, in subsequent foraging sessions, these flowers were gradually positioned further underneath the clear Plexiglas sheet (**Fig. 1C**). Steps I-II did not require any manipulation of the flower itself, with the reward accessible through the extension of the proboscis through the gap beneath the Plexiglas sheet: i.e., the flower is not necessarily manipulated towards the individual and can remain in its initial position. However, the bees learn to reach beneath the gap to obtain the reward. By Step III, the flower *does* need to be brought towards the bee for the reward to be extracted, but this can be achieved by moving the foam disc. In Step IV, the final step, this is no longer possible: the artificial flower is positioned ∼2.0 cm under the Plexiglas, meaning bees must manipulate the string with either their legs or mandibles to pull the flower out from underneath. For Steps I–III, each bee was given 5 foraging sessions, each lasting 5 minutes. If the bee failed to successfully extract a reward within 5 minutes, the artificial flower was moved back to the previous step in the training sequence, requiring the bee to repeat that step in its entirety (i.e., to Step II if the bee had reached Step III). This was done to ensure that the bee could successfully obtain a reward in each session, which maintained their motivation to continue training. After successfully obtaining a reward in Step III, the bees moved on to a final Step IV, where the reward could only be accessed by directly manipulating the string. Once a bee successfully obtained the reward through string pulling for the first time in Step IV, it was tested ten more times with ten artificial flowers. If the bee successfully demonstrated string pulling in all ten tests, it was considered fully trained and became a demonstrator. For examples of successful string pulling, see Supplementary Videos 1-3.

*Phase I*. To investigate the spread of string-pulling as a cultural tradition, we adopted an ‘open-diffusion’ paradigm (Whiten and Mesoudi, 2008). Here, all individuals within a population are allowed to freely interact with their conspecifics and the necessary materials to perform a target behaviour, without encouragement or hindrance from experimenters. Similar open diffusion experiments have been successfully conducted in bumblebees previously, though with a different task (Bridges *et al*., 2023). Control colonies underwent the same procedures as the experimental colonies, but lacked trained demonstrators.

During *Phase I*, open diffusion experiments were conducted every day for five consecutive days. With a single 3 h experimental session conducted per day, bees had a total of 15 h exposure to the string-pulling task during this phase. Before each experimental session, bees were given a 1 h ‘refreshment’ pretraining period, where they were allowed to forage on stringless, randomly-placed, readily-accessible artificial flowers (n=8, each containing 50 μl 50% w/w sucrose solution). This ensured that the association between the artificial flowers and the reward was maintained, and also served to raise individual foraging motivation before the experimental session, which began immediately after pretraining. Here, bees were presented with n=8 artificial flowers positioned underneath the Plexiglas sheet as in Step IV of demonstrator training, with only the strings accessible, for a period of 1 h. Flowers were evenly spaced (15.0 cm apart), with four on each side of the arena: this limited undesirable mass aggregations of foraging individuals in any one spot. Flowers were continuously refilled when depleted, using metal wire hooks to pull them through the gap in the side of the arena (**Fig. 1A**). After the experimental period was over, the artificial flowers were removed, all bees were returned to the nest box, and the arena interior, artificial flowers and strings were cleaned with 70% ethanol. This was done to reduce any possible confounding influence from olfactory cues that might have been left behind (‘residual traces’; see Sherry and Galef, 1984).

*Phase II*. The *Phase II* open diffusion experiments occurred immediately after *Phase I* and followed the same protocol, but the divider was removed from the flight arena to allow two colonies (one *Phase I* experimental colony and one naïve secondary colony) to forage together on n=12 artificial flowers, each containing 50 μl 50% w/w sucrose solution (**Fig. 1A**). *Phase II* continued for 5 days, with a single 3 h experimental session per day. This resulted in a total of 15 h exposure to the string-pulling task during *Phase II*, and 30 h when combined with *Phase I*. After each *Phase II* experimental session ended, bees were sorted and returned to their original nest boxes.

### Video analysis

All observations were recorded using Handycam HDR-CX 190E cameras (Sony, Tokyo, Japan) positioned on the top of the transparent flight arena lid. A total of four cameras were used per diffusion experiment, with each field of view capturing two artificial flowers (**Fig. 1A**). Thus, *Phase I* elicited 4 x 15 h video for each experimental colony (180 h total) and 4 x 15 h video of each control colony (180 h total); with a further 4 x 15 h video being obtained during each *Phase II* experiment (180 h total). Thus, 540 h of video footage in total was obtained for analysis. We used BORIS software (Friard and Gamba, 2016) to record observed string-pulling behaviours as point events. As each bee was tagged, we were able to assign each recorded string-pulling occurrence to the ID of the exact individual who performed it, building a picture of how each bee interacted with the string-pulling task over time. For the extraction of the flower to be assigned to a specific ID, the bee had to either fully extract the flower, or initiate/finalise the extraction by clearly manipulating the string for at least 50% of the required length to obtain the reward while surrounded by conspecifics. If the extraction could not be clearly attributed to an individual bee (e.g., multiple bees manipulated the string simultaneously, or the flower was extracted by squeezing under the table rather than directly manipulating the string with mandibles and legs; see Supplementary Video 4 for an example), the event was left unassigned and excluded from the final analysis. The criterion used to determine whether an individual had learned was the same as our previous work (Bridges *et al*., 2023): once a bee had performed string-pulling twice, it was considered to have learned the behaviour.

### Statistical analysis

Data were plotted and analysed using RStudio v. 3.2.2 (Johnson and Omland, 2004) (R Foundation for Statistical Computing, Vienna, Austria). We conducted generalized linear mixed models (GLMMs) with model selection based on AIC comparisons. Specifically, we considered the model with the lowest AIC score the best model, i.e., the model that provides a satisfactory explanation of the variation in the data (Johnson and Omland, 2004). Following accepted convention, models with an AIC difference of less than 2 units were considered not significantly better than the model it is being compared to (Burnham and Anderson, 2004), thus the simpler model was selected. Model diagnostics were performed using the DHARMa package (Hartig, 2016) to assess and validate model fit. All final model summaries were reported with ANOVA using the car package (Fox, Weisberg and Price, 2001). Post-hoc tests and pairwise comparisons were performed using the emmeans package (Lenth, 2017) and adjusted using Bonferroni corrections when needed. To analyse learning across days (both *Phase I* and *Phase II*, **Fig. 2A-B**, **Fig. 3A-B**), we applied Poisson or negative binomial generalized linear mixed models (GLMMs) depending on model fit. The response variables were the cumulative number of learners or the total number of pulls, with fixed effects for observation day, demonstrator presence (experimental or control for *Phase I*), and colony type (primary or secondary for *Phase II*). Random effects were included for the individual colonies tested.

**Figure 2.**
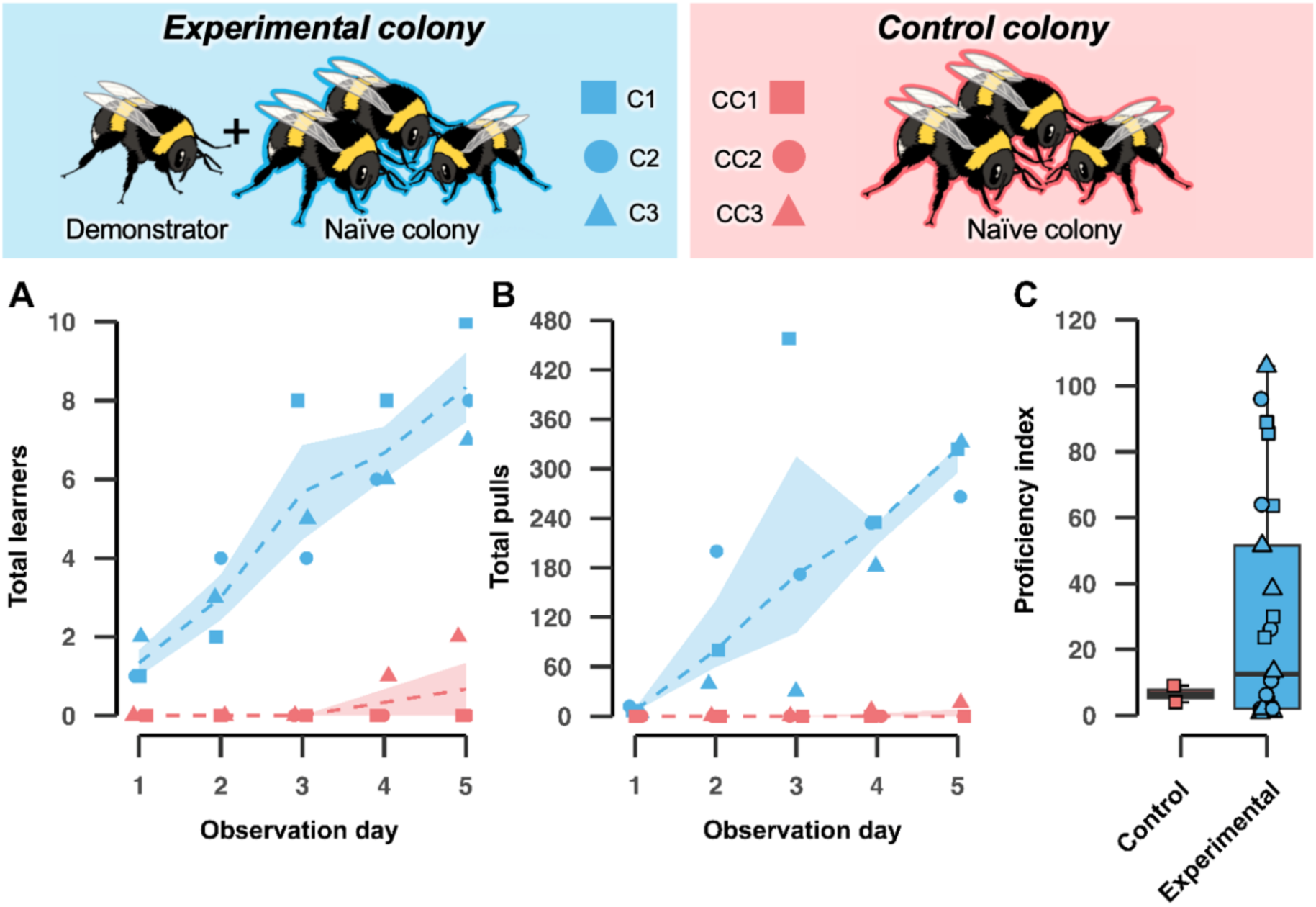
Social learning drives the acquisition and proficiency of the string-pulling technique in bumblebees. **(A)** The number of learners and **(B)** pulling events both steadily increased in experimental colonies (blue), whereas control colonies (red) showed minimal learning. The dashed lines depict the average number of learners within the group (experimental or control) across days, while shaded areas show the interquartile range for experimental (blue) and control (red) colonies. **(C)** The proficiency index, calculated as the total pulling occurrences/number of days spent as a learner, compares individual performance between the groups. Although not statistically significant (GLMM, *χ²* = 1.51, p = 0.09), bees in experimental colonies (blue) exhibited greater overall proficiency in string-pulling behaviour compared to those in control colonies (red).

**Figure 3.**
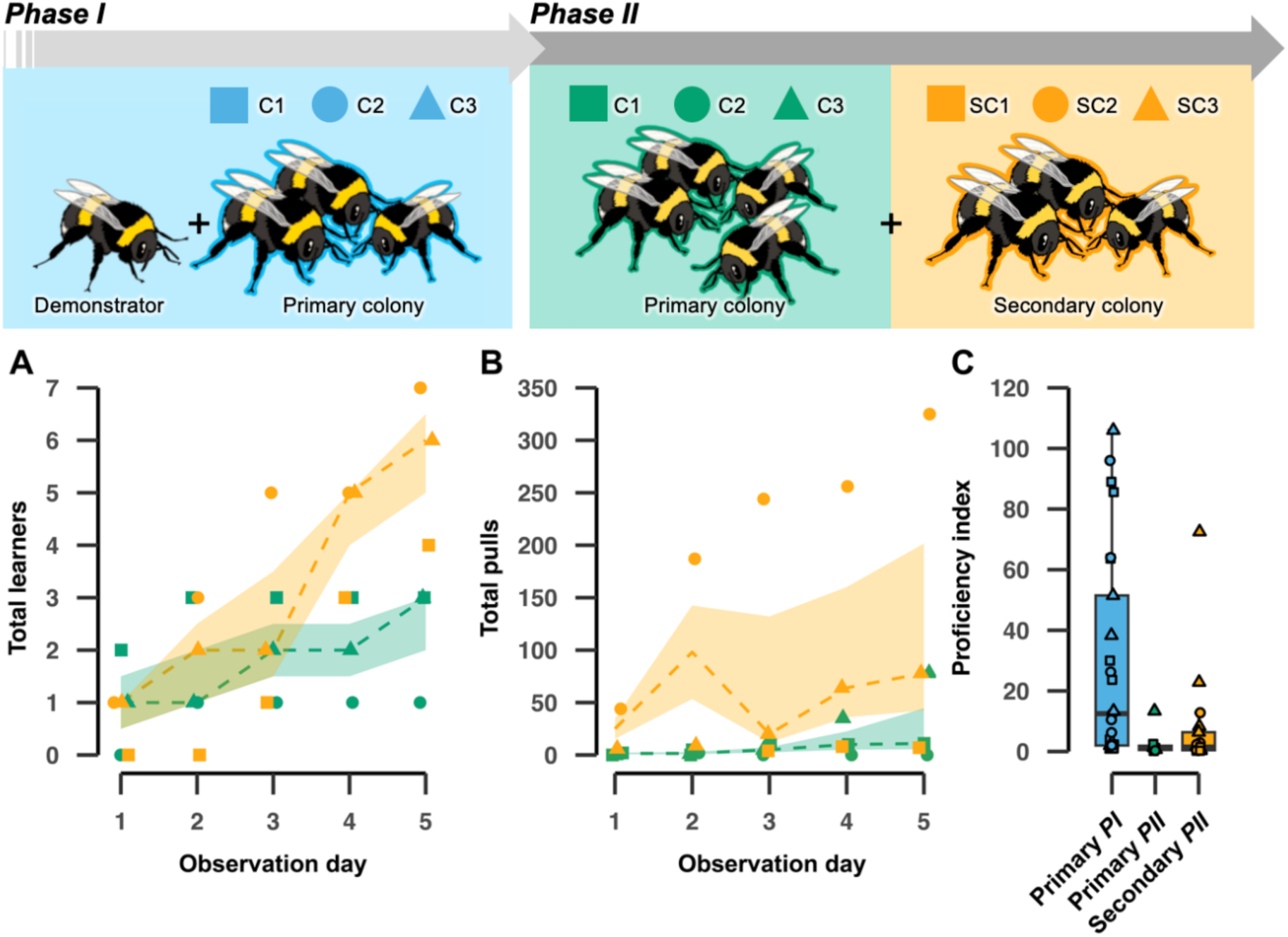
Social learning facilitates the spread of string-pulling behaviour across unrelated colonies. **(A)** The number of new learners (bees that newly reached the learning criterion during *Phase II*) and **(B)** pulling events recorded during *Phase II* increased steadily in both primary (green) and secondary (yellow) colonies, with a faster rate of learning observed in secondary colonies from Day 3 onward. The increase in the number of string-pulling events over time was also more pronounced in secondary colonies (yellow) compared with primary colonies (green) over the *Phase II* diffusion period. The dashed lines depict the group’s average number of learners (A) or pulling events (B) across observation days, with shaded areas showing the interquartile range for primary (green) and secondary (yellow) colonies. **(C)** The proficiency index, calculated as the total pulling occurrences/number of days spent as a learner, compares individual performance across three groups: primary colonies in *Phase I* (blue), primary colonies in *Phase II* (green), and secondary colonies in *Phase II* (yellow).

For overall learning proficiency analysis (**Fig. 2C**, **Fig. 3C**), we performed a Gamma GLMM with a log link. The proficiency index was calculated as previously described (Bridges *et al*., 2023) and defined as the total pulling occurrences divided by the number of days spent as a learner, adjusted for the day the learning criterion was met. When comparing proficiency between experimental and control colonies in *Phase I*, the models included fixed effects for the presence of a demonstrator, with colony as a random factor. For comparisons of learning proficiency between primary and secondary colonies, fixed effects of colony type (*primary Phase I*, *primary Phase II* and *secondary Phase II*) were included, with colony as a random factor.

Individual bee proficiency across the *Phase I* and *II* diffusion experiments (calculated as the ratio of an individual’s total string-pulling occurrences on a given day to the total number of string-pulling occurrences by the entire experimental population on that day) was analysed using Beta family GLMMs. The number of “pullers” (total number of bees demonstrating string pulling on a given day), observation day, colony type (primary or secondary), and learning timing (three level factor, with “early learners” meeting criterion on days 1–3, “mid learners” on days 4–6, and “late learners” on days 7–10) were included as fixed factors. To account for individual variability, bee ID was included as a random factor. Significant interactions were further explored using post-hoc pairwise comparisons of the slopes. P<0.05 was considered to indicate a statistically significant difference.

## Results

### Social learning drives the acquisition of and proficiency with a novel foraging technique in bumblebees

By the end of the *Phase I* open diffusion experiment, untrained bees from all three experimental colonies had successfully acquired string pulling (colony C1, n=10, C2, n=8 and C3, n=7). Notably, while no learners emerged in the control colonies CC1 or CC2, untrained bees (n=2) from control colony CC3 *did* learn string-pulling; despite having no trained demonstrator to model (**Fig. 2A**). This finding was consistent with previous work on string-pulling, which suggested that some bees can learn to pull strings without a demonstrator (Alem *et al*., 2016). However, these learners were fewer in number than those from the experimental colonies, and appeared to begin meeting the learning criterion later (on day 4, compared with as early as day 1 in the experimental colonies; **Fig. 2A**). To assess whether the number of new learners varied significantly across days between the experimental and control colonies, we fitted a Poisson generalised linear mixed-effects model. The best-fit model included observation day and the presence of a demonstrator as main effects, but no interaction between these (see **Supplementary Table S1A.** for model selection details). As anticipated, both factors significantly influenced the cumulative number of naïve bees learning string-pulling over the five days (**Fig. 2A**). Specifically, the number of learners significantly increased as the days of observation progressed (*χ²* = 20.20, *p* < 0.0001) and in the presence of a demonstrator (*χ²* = 29.89, *p* < 0.0001; **Supplementary Table S1B**), underscoring the importance of both temporal dynamics and social learning in the acquisition of the behaviour.

Next, we investigated whether the number of string-pulling instances by learners changed over the course of the diffusion experiment, and whether this change differed between control and experimental colonies: in short, did learners become more proficient at string-pulling with time, and did their proficiency increase more rapidly with a demonstrator present? The best-fitting model included both observation day and the presence of a demonstrator as main effects, with no interaction between these (see **Supplementary Table S2A** for model selection details). As with the total number of learners, the number of pulling events steadily increased across the observation days in both groups (*χ²* = 28.08, *p* < 0.0001; **Supplementary Table S2B**). However, the presence of a demonstrator significantly increased the overall number of pulling instances compared with the control (*χ²* = 33.26, *p* < 0.0001; **Fig. 2B; Supplementary Table S2B**). These findings suggest that while some bees are capable of innovating and learning the task in the absence of a demonstrator, having a demonstrator present led to a substantial increase in both the number of bees acquiring the new technique (**Fig. 2A**) and the frequency with which they performed it (**Fig. 2B**).

Interestingly, despite this latter result, there was no significant difference in terms of *proficiency* between the experimental and control learners (*χ²* = 2.72, *p* = 0.09, **Supplementary Table S3**). As described previously (Bridges *et al*., 2023), since different learners acquired string-pulling on different days, to compare their proficiency we used an index: the total pulling occurrences divided by the number of days spent as a learner. The lack of significance here may be due to the small sample size for the control learners (n=2; with an average proficiency index of 6.5 ± 2.5; median ± IQR, **Fig. 2C**). In contrast, the experimental group (n=25) exhibited substantial variation in proficiency levels, with an average proficiency index of 12.5 ± 49.5 (median ± IQR, **Fig. 2C**). While the highest recorded proficiency index was 106, a notable portion of experimental learners (9 out of 25) had a proficiency index <4: lower than the smallest value observed in the control group. The control learners both remained at the lower end of this scale, with proficiency indexes of 4 and 9. Thus, despite the lack of significance in individual proficiency, our overall results did broadly align with those of our previous open diffusion experiments (Bridges *et al*., 2023), which suggested that while some bees have the capacity to innovate a novel behaviour, it is the social influence of a knowledgeable conspecific that allows it to become established in a group as a local adaptation.

### Social learning also facilitates the spread of string-pulling behaviour between unrelated colonies

After confirming that string-pulling acquisition and proficiency within a single colony was highly dependent on the presence of a demonstrator in our open diffusion paradigm, we next examined whether this learned behaviour could continue to spread beyond this initial colony. Specifically, we sought to determine whether string-pulling could be transmitted to secondary colonies of genetically unrelated naïve bees; continuing the learning process across colony boundaries. At the end of *Phase I*, 10, 8, and 7 naïve bees in the experimental colonies C1, C2 and C3, respectively, had successfully met the learning criterion. In *Phase II*, we continued the open diffusion experiments by introducing a new secondary colony to each primary colony, and recorded the number of new bees that acquired string-pulling in both colonies.

As before, we assessed whether the number of new learners continued to increase over *Phase II,* and whether this rate of increase differed between primary colonies (those exposed to five days of open diffusion previously, during *Phase I*) and secondary colonies (newly-introduced and naïve to the string-pulling technique at the start of *Phase II*). Indeed, bees from all three secondary colonies learned to string-pull during *Phase II* (SC1, n=4, SC2, n=7 and SC3, n=6; Fig. 3A). When controlling for the random effect of colony ID, the best-fitting model included fixed effects for observation day and colony type (primary or secondary), along with their interaction (see **Supplementary Table S4.1A** for model selection details). Our analysis revealed a significant increase in the number of new learners across the five days in both the primary and secondary colonies during *Phase II* (*F* = 26.98, *p* < 0.0001; **Supplementary Table S4.1B**). While the fixed effect of colony type was not statistically significant (*F* = 1.59, *p* = 0.276), the significant interaction between colony type and observation day (*F* = 10.88, *p* = 0.00019) indicates that learning rates varied between primary and secondary colonies over time. Post hoc analyses showed no significant differences in the number of new learners between the two colony types during Days 1 to 3 (**Fig. 3A**; **Supplementary Table S4.1B**). However, starting on Day 4, a higher number of new learners emerged from the secondary colonies, with the difference becoming statistically significant on Day 5 (estimate ± s.e. = - 3.33 ± 1.04, *t* = -3.19, *p* = 0.020, **Supplementary Table S4.2**). This pattern suggests that secondary colonies displayed a faster learning rate during the later days of *Phase II*. This may have been because by this stage, most of the available motivated foragers from the primary colony had already learned string-pulling.

Next, we assessed the number of string-pulling occurrences, using the total number of pulls by untrained bees as the response variable (see **Supplementary Table S5A**, for model selection details). This analysis revealed that pulling events by untrained bees also increased significantly with observation day (*χ²* = 37.28, *p* < 0.0001), with learners from secondary colonies performing string-pulling significantly more often than those from the primary colonies during *Phase II* (*χ²* = 5.01, p = 0.025); **Fig. 3B**; **Supplementary Table S5B**). Despite pulling more frequently, however, post hoc analyses showed that by the end of *Phase II*, the proficiency of learners from secondary colonies was not significantly different from those in primary colonies that learned during either *Phase II* (*p* = 0.647) or *Phase I* (*p* = 0.100; **Fig. 3C**, **Supplementary Table S6.2**), when comparing their proficiency indexes. Interestingly, there was a significant decrease in proficiency in learners from primary colonies in *Phase II* compared to learners from primary colonies in *Phase I* (*p* = 0.0002; **Supplementary Table S6.2**), indicating a substantial decline in learning outcomes in primary colonies between phases.

Taken together, these findings demonstrated that, once established, string-pulling can spread rapidly from experienced bees in the primary colonies to naïve bees in secondary colonies. Social learning continued to drive the diffusion of string-pulling across colony boundaries: in theory, this could continue for as long as there are foragers available to learn.

### The number of knowledgeable bees present may have impacted the acquisition of string-pulling by new bees

The main difference between the conditions naïve bees were exposed to in *Phase I* and *II* was the number of knowledgeable bees present at the start. In *Phase I*, the diffusion process began with the introduction of a single trained demonstrator. However, *Phase II* began with multiple knowledgeable foragers present from the start. To investigate the impact of this on the diffusion of string-pulling, we directly compared learner accumulation and string-pulling performance between primary colonies during the five days of *Phase I*, and secondary colonies during the five days of *Phase II*.

The best-fitting model comparing the number of cumulative learners between these groups included observation day, colony type (primary vs. secondary), without interaction (**Supplementary Table S7)**. As before, observation day was a highly significant predictor of learner number, with more learners having emerged by later experimental days (*χ²* = 32.23, *p* < 0.0001). Colony type, however, was also highly significant (*χ²* = 7.34, *p* = 0.007), indicating that the difference in the number of learners between primary *Phase I* and secondary *Phase II* remained constant across days.

For the number of pulling events, the best-fitting model included observation day, colony type, and their interaction (**Supplementary Table S8.1**). In both colony types, there was a significant increase in pulling events recorded over the diffusion days (*χ²* = 530.91, *p <* 0.0001). Colony type was not significant as a fixed factor; however, the significant interaction between colony type and observation day indicates that the effect of group is dependent on the day (*χ²* = 52.84, *p* < 0.0001). Indeed, post hoc analysis revealed a trend in which primary *Phase I* colonies consistently exhibited more pulling events compared to secondary *Phase II* colonies (positive estimates for all days; see **Supplementary Table S8.2**). While these differences were not significant at the beginning of diffusion (days 1 and 2: estimate ± s.e. = 0.09 ± 0.94, *z* = 0.09, *p* = 0.925, and 1.35 ± 0.91, *z* = 1.49, *p* = 0.135, respectively), they became statistically significant by day 3 (estimate ± s.e. = 1.79 ± 0.90, *z* = 1.98, *p* = 0.048). On subsequent days, the differences approached significance (day 4: estimate ± s.e. = 1.57 ± 0.90, *z* = 1.74, *p* = 0.082; day 5: estimate ± s.e. = 1.70 ± 0.90, *z* = 1.88, *p* = 0.06) but did not reach statistical significance after Bonferroni adjustment. These results suggest that differences in pulling behaviour between colony types became more pronounced over time, particularly starting from observation day 3, with primary colonies generally showing a stronger tendency for more pulling events compared to secondary colonies.

### Restricted resources may limit observer performance in the string-pulling task under open diffusion conditions

The trends we observed between the primary *Phase I* and secondary *Phase II* colonies seem counterintuitive. If the presence of a demonstrator is key to the spread of string-pulling, as seen when comparing experimental colonies to controls, then why would an increased availability of demonstrators be no better, or worse, than having just one? Thus, we investigated whether other factors might be influencing the spread of string-pulling in our experimental set-up.

Many naïve bees (n=24 overall, with n=7 from primary colonies and n=17 from secondary colonies) successfully reached the learning criterion during *Phase II*. As aforementioned, while there was no significant difference between the proficiency of learners from the primary and the secondary colonies, there was still a decline in the overall proficiency of learners from *Phase I to Phase II* (**Fig. 3C**). This difference was most notable among learners from the primary colonies, with a significant decline from *Phase I* to *II* (*p* = 0.0002; **Fig. 3C; Supplementary Table S6.2**). One explanation might be that the bees from the primary colonies, who learned during *Phase I*, might be less active due to, for example, age. However, this alone did not explain the downward trend among learners from the *Phase II* secondary colonies compared with those from *Phase I* primary colonies (**Fig. 3C**).

Another possible explanation is that, as the number of knowledgeable bees present in the flight arena increased, the limited availability of artificial flowers for string pulling (with a maximum of 12 flowers available at any given time) resulted in reduced opportunities for new bees to pull strings and demonstrate their learning compared with at the start of the experiment (**Fig. 4**). To explore this further, we investigated how the number of pullers active on a given day (*no. of pullers*) influenced individual pulling efficiency on that day. We also considered whether colony type (*introduction*: primary or secondary), observation day (*day*) and when the bee met the learning criterion (*criterion met*: early learners, on days 1-3; mid learners, on days 4-6; late learners, on days 7-10) had an effect on daily performance. The best-fitting model, which controlled for individual variation as a random effect, included fixed effects for the number of pullers each day (*no. of pullers*), learning time (*criterion met*), colony type (*introduction phase*), and observation day (*day*), as well as an interaction between the number of pullers and observation day (see **Supplementary Table S9.1A** for model selection details).

**Figure 4.**
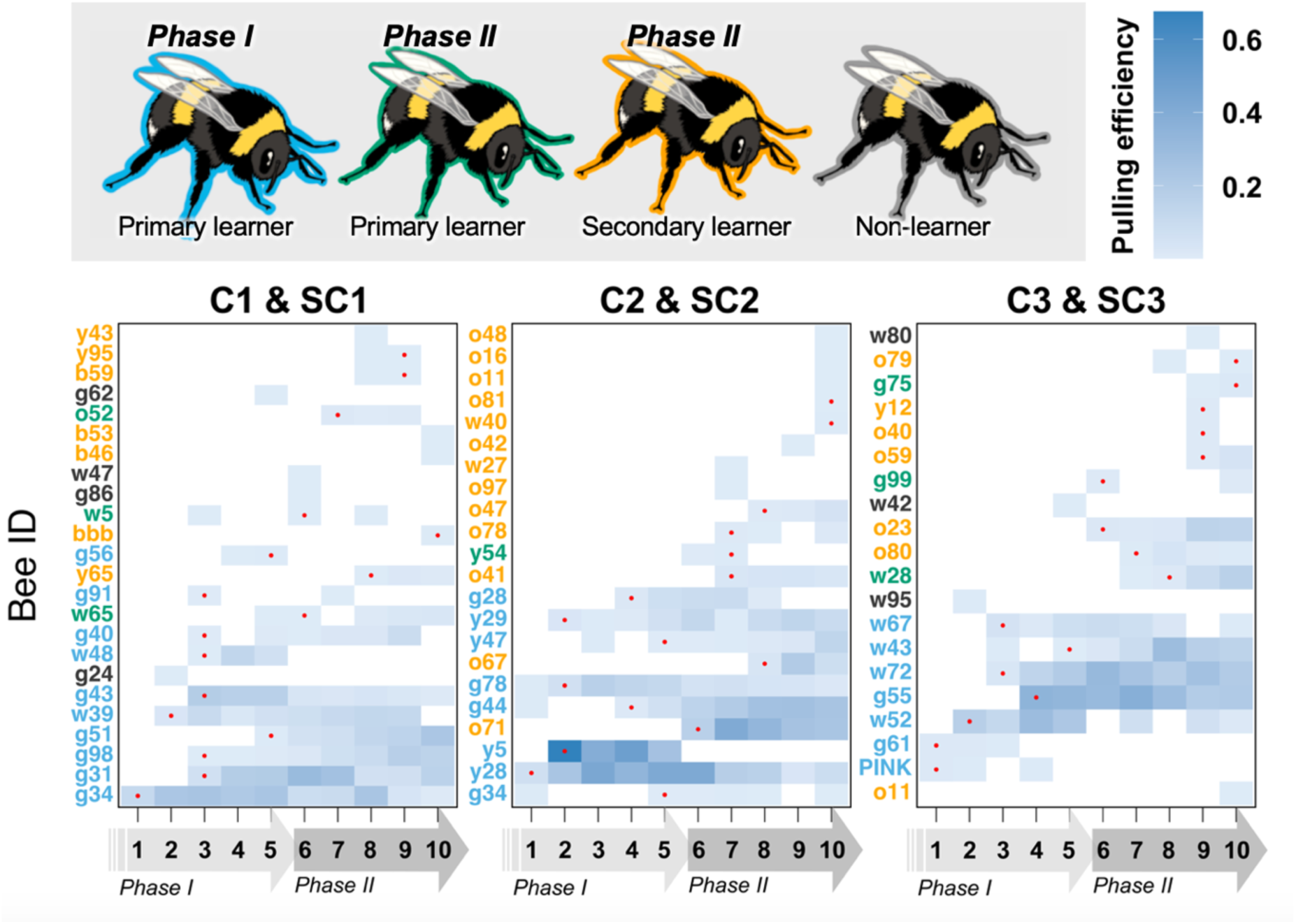
Restricted resources may limit observer string-pulling performance under open diffusion conditions. The raster plots illustrate individual pulling efficiency (total pulling occurrences for each bee adjusted for the total occurrences across all bees for a given day) across the observational days, for three experimental colonies during *Phase I* and *Phase II*: *Experimental Colony 1* (primary colony C1 with a seeded demonstrator and secondary introduced naïve colony SC1), *Experimental Colony 2* (primary colony C2 with a seeded demonstrator and secondary introduced naïve colony SC2), and *Experimental Colony 3* (primary colony C3 with a seeded demonstrator and secondary introduced naïve colony SC3). Each row represents a single bee (with bee IDs from primary and secondary colonies shown in green and yellow, respectively). The level of shading indicates the proportion of total pulls attributed to each bee on a given day, with darker shading representing a higher number of pulls. Red dots indicate the day on which each bee met the learning criterion (successfully pulling the string for the second time).

The analysis revealed that observation day did not have a significant effect on performance (*χ2* = 11.08, *p* = 0.270), indicating that performance changes are not driven simply by experimental progression but are rather impacted by other factors. Specifically, colony type (*introduction phase*) significantly influenced performance (*χ2* = 5.48, *p* = 0.019), with bees from primary colonies outperforming those from secondary colonies (estimate ± s.e. = 0.694 ± 0.296, *p* = 0.019). Performance also varied significantly based on when a bee met the learning criterion (*χ2* = 13.39, *p* = 0.001). Post-hoc paired comparisons revealed that early learners had significantly better overall performance than mid learners (estimate ± s.e. = 1.59 ± 0.59, *p* = 0.019) and late learners (estimate ± s.e. = 0.74 ± 0.28, *p* = 0.021). At the same time, mid learners and late learners did not differ (estimate ± s.e. = -0.85 ± 0.60, *p* = 0.335).

The number of pullers on a given day (*no. of pullers*) did not directly influence performance, though it showed a marginally significant effect (*χ2* = 3.56, *p* = 0.059). However, there was a significant interaction between the number of pullers and when bees met the learning criterion (*χ2* = 7.30, *p* = 0.026). Indeed, post-hoc analyses revealed that early learners showed a strong trend toward declining performance as the number of learners increased (estimate ± s.e. = -0.07 ± 0.04, *p* = 0.059). In contrast, neither mid learners (estimate ± s.e. = 0.185 ± 0.145, p = 0.205) nor late learners (estimate ± s.e. = 0.03 ± 0.04, p = 0.496) exhibited a significant improvement in performance as the number of learners increased. This stability may be due to their lower baseline performance than the early learners, being unable to rack up large numbers of pulls in any one observation session due to increased competition from other learners. In short, our results do not necessarily suggest that bees that learned later were *incapable* of pulling as many strings as those that learned earlier: rather, they did not get the chance to do so due to the limitations of our experimental design.

## Discussion

The results of the present study represent a clear confirmation of the potential for local behavioural adaptations to emerge among groups of bumblebees, if opportunity arises. The spread of string-pulling from a single, experimenter-trained individual to multiple untrained observers, who then repeatedly performed the behaviour themselves, fulfils the definition of culture as used in animals (Fragaszy and Perry, 2003). A similar phenomenon was observed in previous work by our group (Bridges *et al*., 2023), using a different non-natural foraging paradigm: a puzzle-box that had to be opened by rotating a lid. However, the present study builds on this further by demonstrating the spread of string-pulling between *unrelated* colonies of bumblebees. Once string-pulling was established in our primary experimental colonies, it readily spread through newly-introduced secondary colonies. This confirms, at least in theory, that socially-learned behaviour could spread beyond localised, related groups of bumblebees should it ever arise in the field.

Further analysis of our results provided additional insights about the spread of string-pulling in our open diffusion paradigm. First, while the number of new learners from both primary and secondary colonies significantly increased during *Phase II*, towards the end of this phase significantly more learners emerged from the secondary colonies. This may simply have been due to the pool of potential learners available within the primary colonies being exhausted by that point. *B. terrestris* colonies can grow to contain more than two hundred workers (Sladen, 1912), but not all of these bees act as foragers at any one time. Although they do not show the strict age-dependent polyethism of honeybee workers, younger bumblebees tend to perform tasks within the nest and switch to foraging later on, although some individuals never make the switch (Hoffer, 1882; Sladen, 1912; Goulson, 2010). It is also possible that some of these foragers will never learn to string pull. Lack of motivation, neophobia, or limitations in terms of cognitive capacity may reasonably be posited to prevent some individuals ever acquiring such behaviour. Any of these factors might limit the number of potential string-pullers present in a single colony at any one time. However, the secondary colonies provided an additional pool of potential learners and allowed string-pulling to continue to spread beyond these potential limitations.

There was also limited evidence to suggest that the learners from the secondary colonies were less proficient than those from the primary colonies. There was no significant difference between the overall proficiency of learners from primary and secondary colonies (Fig. 3C; Supplementary Table S6.2), although there was a decline in overall learner proficiency between *Phase I* and *II*. Particularly in the later stages of the experiment, competition over the limited number of flowers available may have prevented naïve bees from having the chance to learn, or to demonstrate their proficiency. As the number of knowledgeable bees increased, individual proficiency appeared to decrease, but the starkest difference in performance was found among “early” learners – those that first acquired string-pulling in the early days of the experiment. These early learners, facing less competition for flowers, were able to rack up large numbers of pulls without being inhibited by other knowledgeable bees at the start of the experiment, setting a higher “baseline” number of pulls for themselves that then had to decrease. It is also conceivable that this early chance to string-pull many times served to set these learners up better than their mid-to-late-learning rivals from the offset, giving them more chances to practice and refine their string-pulling. In cases where multiple bees competed to pull one string, more experienced early learners would conceivably be more likely to take over and pull faster, further restricting mid-to-late learners within the bounds of our set-up.

It is unlikely that the decrease in proficiency was linked to any decay in the integrity of string-pulling transmission over multiple learning events. In colonies C1/SC1 and C3/SC3, the original trained demonstrators from C1 and 3 remained actively pulling throughout *Phase I* and *II*. However, in colony C2, the demonstrator ceased string-pulling after the first day of *Phase I* (**Supplementary Fig. 5**; **Supplementary Table S10**). This meant that all learners from the corresponding secondary colony SC2 were only ever exposed to knowledgeable but untrained bees. Still, learners o71 and o67 from colony SC2 went on to become the most proficient of all learners from secondary colonies, pulling 745 and 183 strings, respectively, during *Phase II.* (**Supplementary Fig. 2**; **Supplementary Table S10**). Taken together, the root cause of this decline in proficiency appears to have been the ‘artificial ceiling’ imposed by the number of strings available for the bees to pull in our experimental set-up, and the consequences of this, rather than string-pulling failing to be properly transmitted across multiple generations of learners.

Our experiments also elicited another result worthy of further discussion. In our control colonies, which had no seeded demonstrators, fewer bees learned than in the matched experimental colonies. However, some *did* learn. A total of two bees from colony CC3 met the learning criterion, with the most proficient of these performing string-pulling 18 times. This has also been observed in previous dyadic experiments on string-pulling (Alem *et al*., 2016), as well as in open diffusion experiments involving a single-step puzzle box (Bridges *et al*., 2023). This does not preclude string-pulling from having spread via social learning within our paradigm: significantly more bees learned in the experimental colonies. However, it serves to highlight yet again the astonishing behavioural flexibility of bumblebees. Repeatedly, across different non-natural foraging problems, bumblebees manage to innovate their own solutions without any demonstration or training – but not *all* bees. Determining what makes these particular individuals innovate, while others fail to do so, would be a fascinating route for future studies. Such innovators may represent “keystone individuals”, acting as a source of novel behaviours or behavioural variants that can later become traditional within a population (Arbilly, 2018).

To conclude, the results of our study confirm that, when given opportunity, bumblebees can innovate novel behaviours, can socially learn these behaviours, and also have the ability to transmit and acquire these behaviours beyond colony limitations. The opportunity to practise and perform a behaviour, which can be inhibited by competition from other knowledgeable individuals when resources are limited, may also be key. However, the question as to whether similar novel, local behavioural adaptations occur among wild bumblebees remains open. Nectar-robbing and its handedness (Goulson *et al*., 2013) may represent one potential example. However, their historical characterisation as “biological automatons” has often resulted in their behaviour being overlooked (Chittka and Wilson, 2019), especially when it is suggestive of so-called “complex” cognition. Yet, with this and previous studies (Alem *et al*., 2016; Bridges *et al*., 2023), we have shown a clear and replicable tendency for bumblebees to develop their own local behavioural adaptations, which spread via social learning, if they are given the opportunity to do so. If they fail to do so in the wild, therefore, this would suggest a lack of need or opportunity rather than any fundamental capacity.

## Acknowledgements

This study was funded by grants from EPSRC (grant no. EP/P006094/1, awarded to L.C., and EP/X019705/1, awarded to M.J.), BBSRC (BB/F012071/1 and BB/X006247/1, awarded to M.J.), Leverhulme (RPG-2024-016, awarded to M.J.) and a QMUL studentship (awarded to A.D.B.).

## Author contributions

**O.P.:** methodology, investigation, formal analysis, visualization, writing (original draft preparation). **A.D.B.:** conceptualization, methodology, supervision, investigation, visualization and writing (original draft preparation, review and editing). **A.K.:** investigation. **M.J.:** funding acquisition, writing (editing). **L.C.:** funding acquisition, conceptualization, supervision, resources, writing (review and editing).

## Competing interests

The authors have no competing interests to declare.

## Supplementary information

**Table S1.** Comparison and summary of Poisson generalised linear mixed-effects models analysing the impact of demonstrator presence and observation day on the cumulative number of learners recorded during *Phase I*.

| A. Selection of best-fit model |  |  |  |  |  |
| --- | --- | --- | --- | --- | --- |
| Dependent variable | Fixed factors | Random factor | df | AIC | ΔAIC |
| Cumulative learners | Demonstrator + Observation day | Colony ID | 4 | 77.22 | 0 |
|  | *Demonstrator \* Observation day* | *Colony ID* | 5 | 77.04 | 0.18 |
| B. Factor effects for selected model |  |  |  |  |  |
| Dependent variable | Fixed factors | | df | $\chi^2$ | P |
| Cumulative learners | (Intercept) |  | 1 | 19.67 | <0.0001* |
|  | Demonstrator |  | 1 | 29.89 | <0.0001* |
|  | Observation day |  | 1 | 20.20 | <0.0001* |

**A.** Two models were fitted to assess cumulative learners in relation to observation days and demonstrator present, accounting for colony variability. The table shows the best-fitting models, detailing fixed and random factors, degrees of freedom (df), and Akaike's Information Criterion (AIC). The model with the lowest AIC was considered best; however, when competing models had a $\Delta AIC < 2$ , they were not deemed significantly better (Burnham and Anderson, 2004), so the simpler model was chosen. The final selected model is indicated in italics. **B.** \*Significant at $P < 0.05$ .

**Table S2.**
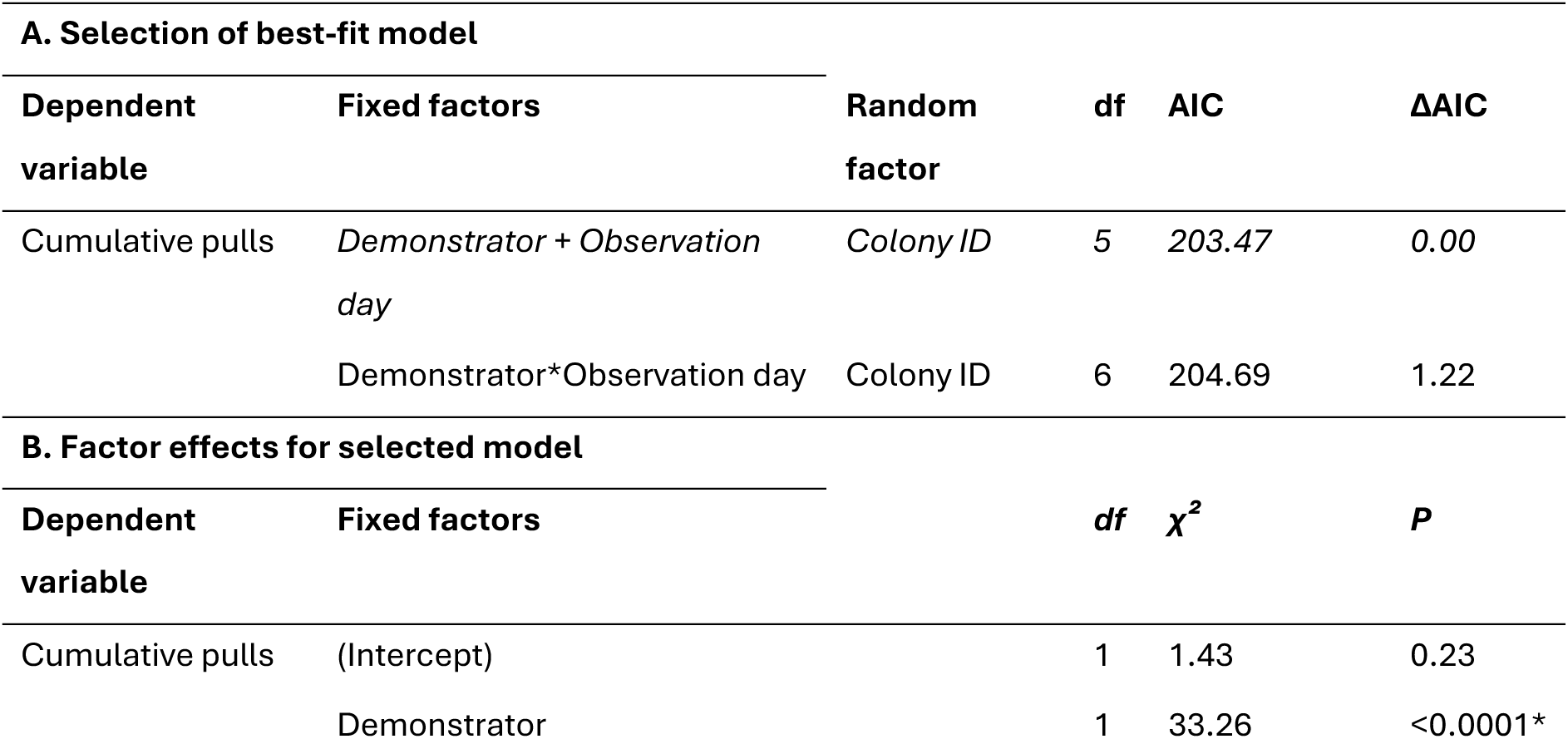
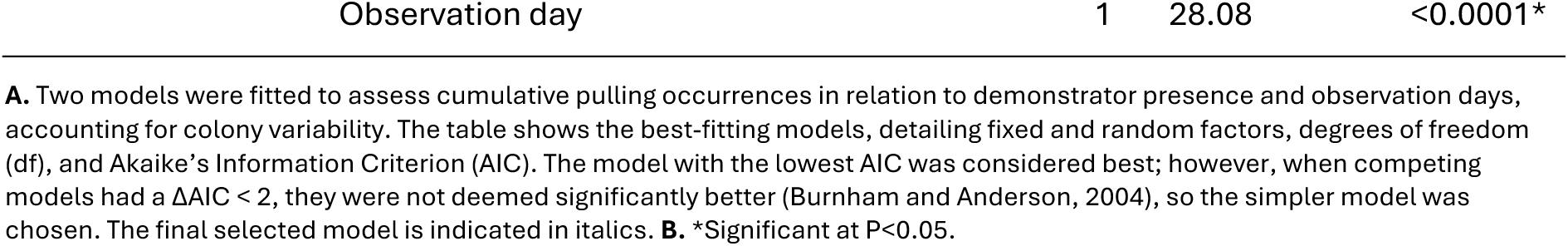
Comparison and summary of negative binomial generalised linear mixed-effects models analysing the impact of demonstrator presence and observation days on the cumulative number of string-pulling occurrences during *Phase I*.

**Table S3.** Summary of the gamma generalized linear mixed-effects model analysing the impact of demonstrator presence vs. absence on overall individual proficiency during *Phase I*.

| A. Best-fit model |  |  |  |  |
| --- | --- | --- | --- | --- |
| Dependent variable | Fixed factors | Random factor |  | AIC |
| Proficiency | Demonstrator | Colony ID |  | 234.42 |
| B. Factor effects for selected model |  |  |  |  |
| Dependent variable | Fixed factors | df | $\chi^2$ | P |
| Proficiency | Demonstrator | 1 | 2.72 | 0.099 |

**A.** A single model was fitted to assess demonstrator presence and the overall individual proficiency, accounting for colony variability. The table shows the fitted model, detailing fixed and random factors, degrees of freedom (df), and Akaike's Information Criterion (AIC). **B.** \*Significant at $P < 0.05$ .

**Table S4.1.**
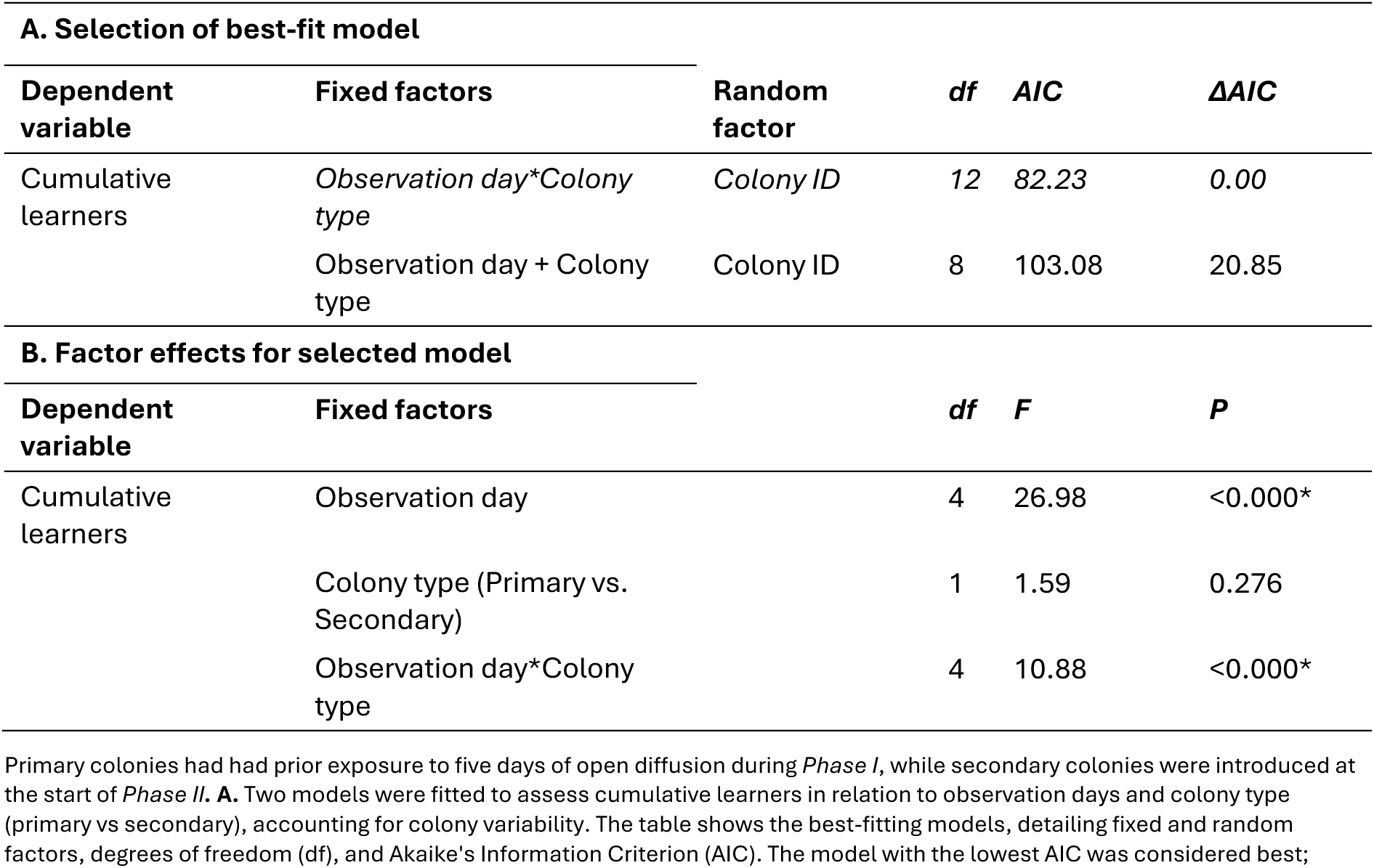
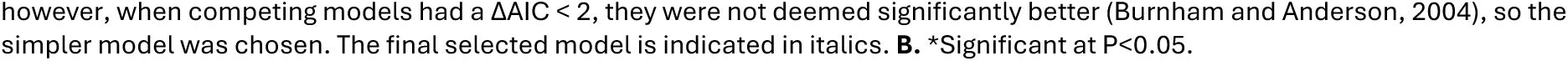
Comparison and summary of best-fit generalized linear mixed-effects models analysing the impact of observation day and colony type (primary vs. secondary) on the cumulative number of learners that emerged during *Phase II*.

**Table S4.2.** Post hoc analysis of the interactions between colony type and observation day, and their impact on the cumulative number of learners recorded during *Phase II*.

| Observation day | Contrast | Estimate | SE | df | t | P |
| --- | --- | --- | --- | --- | --- | --- |
| 1 | Primary vs. Secondary | 0.33 | 1.04 | 5.71 | 0.32 | 0.761 |
| 2 | Primary vs. Secondary | 0.00 | 1.04 | 5.71 | 0.00 | 1.000 |
| 3 | Primary vs. Secondary | -0.67 | 1.04 | 5.71 | -0.64 | 0.548 |
| 4 | Primary vs. Secondary | -2.33 | 1.04 | 5.71 | -2.24 | 0.069 |
| 5 | Primary vs. Secondary | -3.33 | 1.04 | 5.71 | -3.19 | 0.020* |

\*Significant at $P < 0.05$ . The Bonferroni p-value adjustment method used was the R package emmeans. Primary colonies had had prior exposure to five days of open diffusion during *Phase I*, while secondary colonies were introduced at the start of *Phase II*.

**Table S5.**
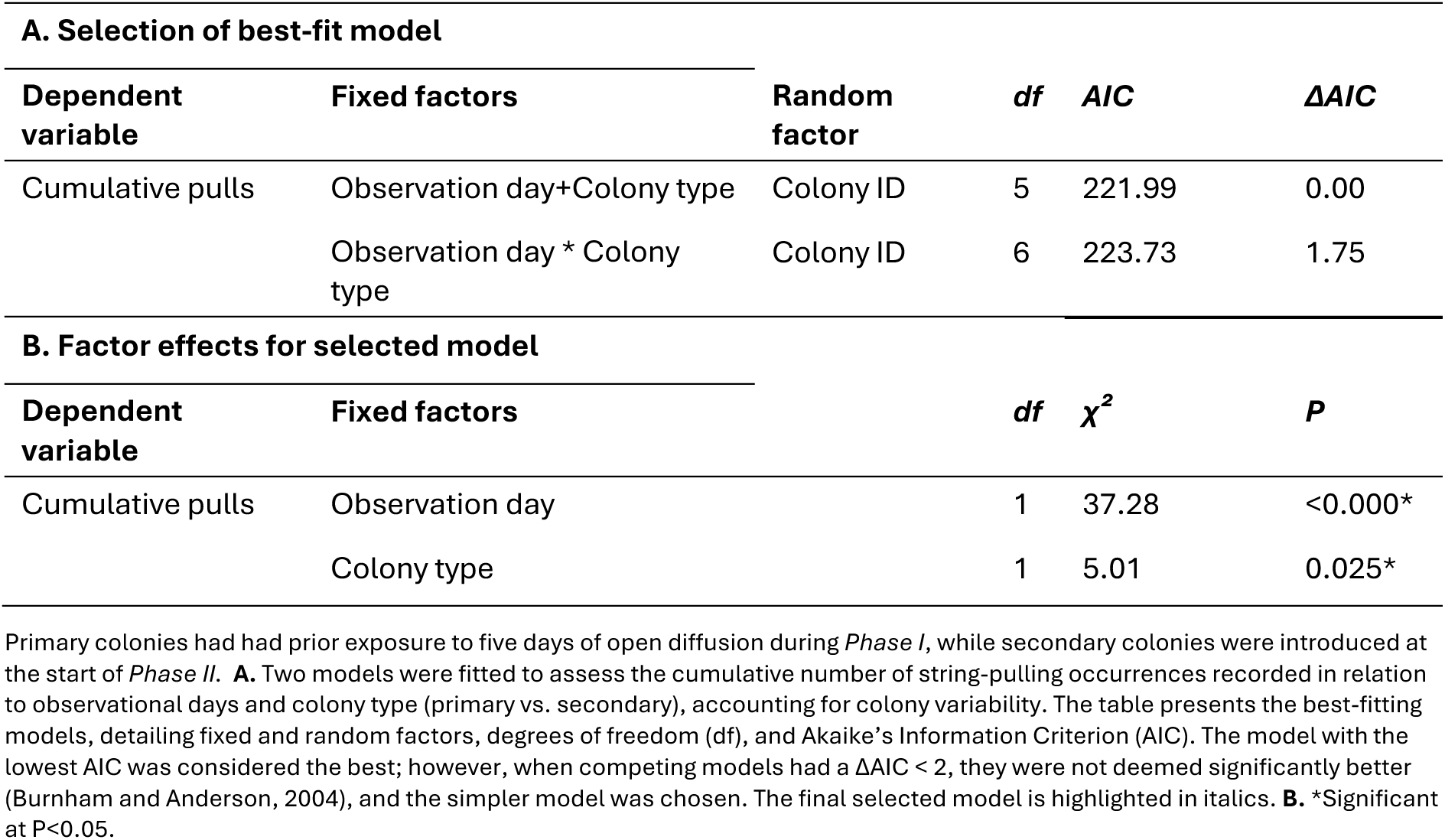
Comparison and summary of best-fit negative binomial generalized mixed models analysing the impact of observation days and colony type (primary vs. secondary) on the cumulative string-pulling occurrences recorded during *Phase II*.

**Table S6.1.**
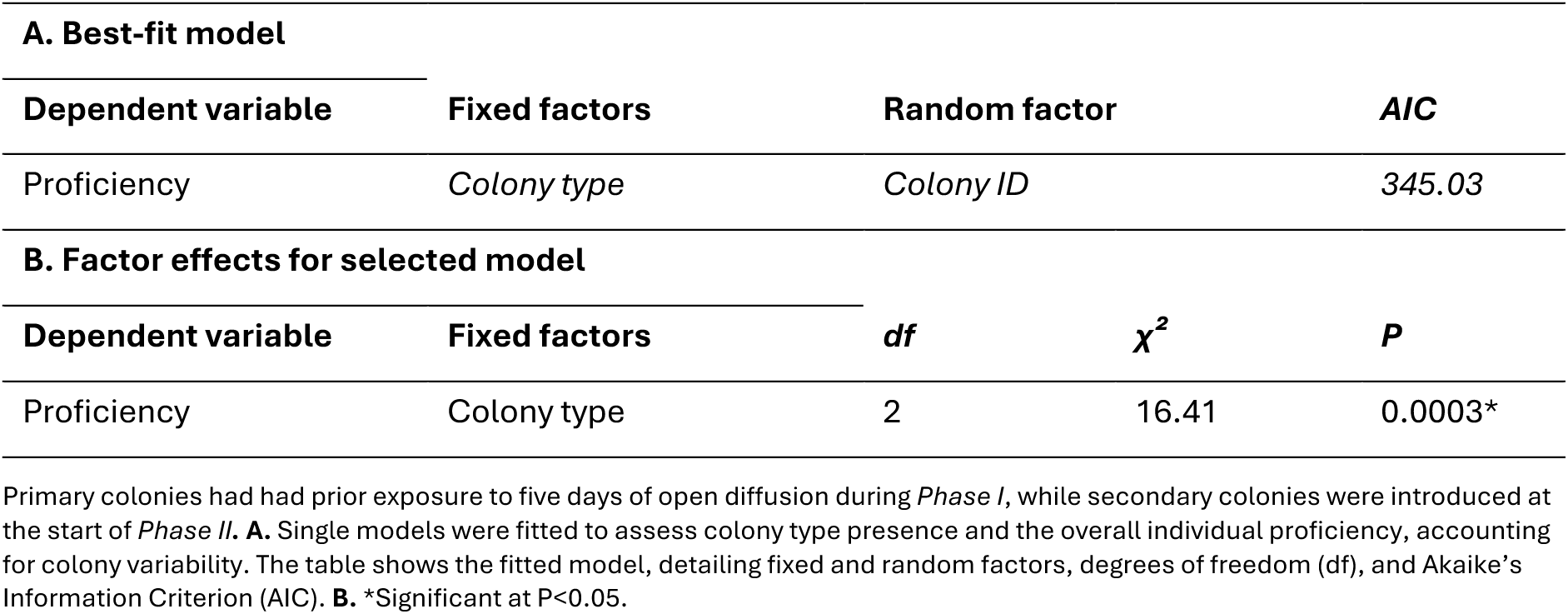
Summary of the best-fit Poisson generalized linear mixed-effects model analyzing the impact of colony type (primary vs. secondary) on the overall individual proficiency of new learners that emerged in *Phase II*.

| A. Best-fit model |  |  |  |  |
| --- | --- | --- | --- | --- |
| Dependent variable | Fixed factors | Random factor |  | AIC |
| Proficiency | Colony type | Colony ID |  | 345.03 |
| B. Factor effects for selected model |  |  |  |  |
| Dependent variable | Fixed factors | df | $\chi^2$ | P |
| Proficiency | Colony type | 2 | 16.41 | 0.0003* |

Primary colonies had had prior exposure to five days of open diffusion during *Phase I*, while secondary colonies were introduced at the start of *Phase II*. **A.** Single models were fitted to assess colony type presence and the overall individual proficiency, accounting for colony variability. The table shows the fitted model, detailing fixed and random factors, degrees of freedom (df), and Akaike's Information Criterion (AIC). **B.** \*Significant at $P < 0.05$ .

**Table S6.2.** Post hoc pairwise comparisons of the overall learning proficiency across learners in primary colonies during *Phase I*, primary colonies during *Phase II*, and secondary colonies during *Phase II* (Bonferroni p-value adjustment method, R package emmeans).

| **Contrast** | **Estimate** | **SE** | **z** | **df** | **P** |
| --- | --- | --- | --- | --- | --- |
| Primary *Phase I* vs Primary *Phase II* | 2.46 | 0.62 | 3.94 | Inf | 0.0002* |
| Primary *Phase I* vs Secondary *Phase II* | 1.70 | 0.83 | 2.05 | Inf | 0.100 |
| Primary *Phase II* vs Secondary *Phase II* | -0.77 | 0.86 | -0.89 | Inf | 0.647 |

\*Significant at $P < 0.05$ . Primary colonies had had prior exposure to five days of open diffusion during *Phase I*, while secondary colonies were introduced at the start of *Phase II*.

**Table S7.** Comparison and summary of best-fit Poisson generalized linear mixed-effects models analyzing the impact of observation day and colony type (primary *Phase I* vs. secondary *Phase II*) on the cumulative number of learners.

| **A. Selection of best-fit model** |  |  |  |  |  |
| --- | --- | --- | --- | --- | --- |
| **Dependent variable** | **Fixed factors** | **Random factor** | **df** | **AIC** | **ΔAIC** |
| Cumulative learners | *Observation day+Colony type* | *Colony ID* | 4 | 113.13 | 0.00 |
|  | *Observation day \* Colony type* | *Colony ID* | 5 | 114.80 | 1.67 |

| B. Factor effects for selected model |  |  |  |  |
| --- | --- | --- | --- | --- |
| Dependent variable | Fixed factors | df | $\chi^2$ | P |
| Cumulative learners | (Intercept) | 1 | 0.64 | 0.42 |
|  | Observation day | 1 | 32.23 | <0.000* |
|  | Colony type | 1 | 7.34 | 0.007* |

A. Two models were fitted to assess cumulative learners in relation to observation days and colony type, accounting for colony variability. The table shows the best-fitting models, detailing fixed and random factors, degrees of freedom (df), and Akaike's Information Criterion (AIC). The model with the lowest AIC was considered best; however, when competing models had a $\Delta AIC < 2$ , they were not deemed significantly better (Burnham and Anderson, 2004), so the simpler model was chosen. The final selected model is indicated in italics. B. \*Significant at $P < 0.05$ .

**Table S8.1.**
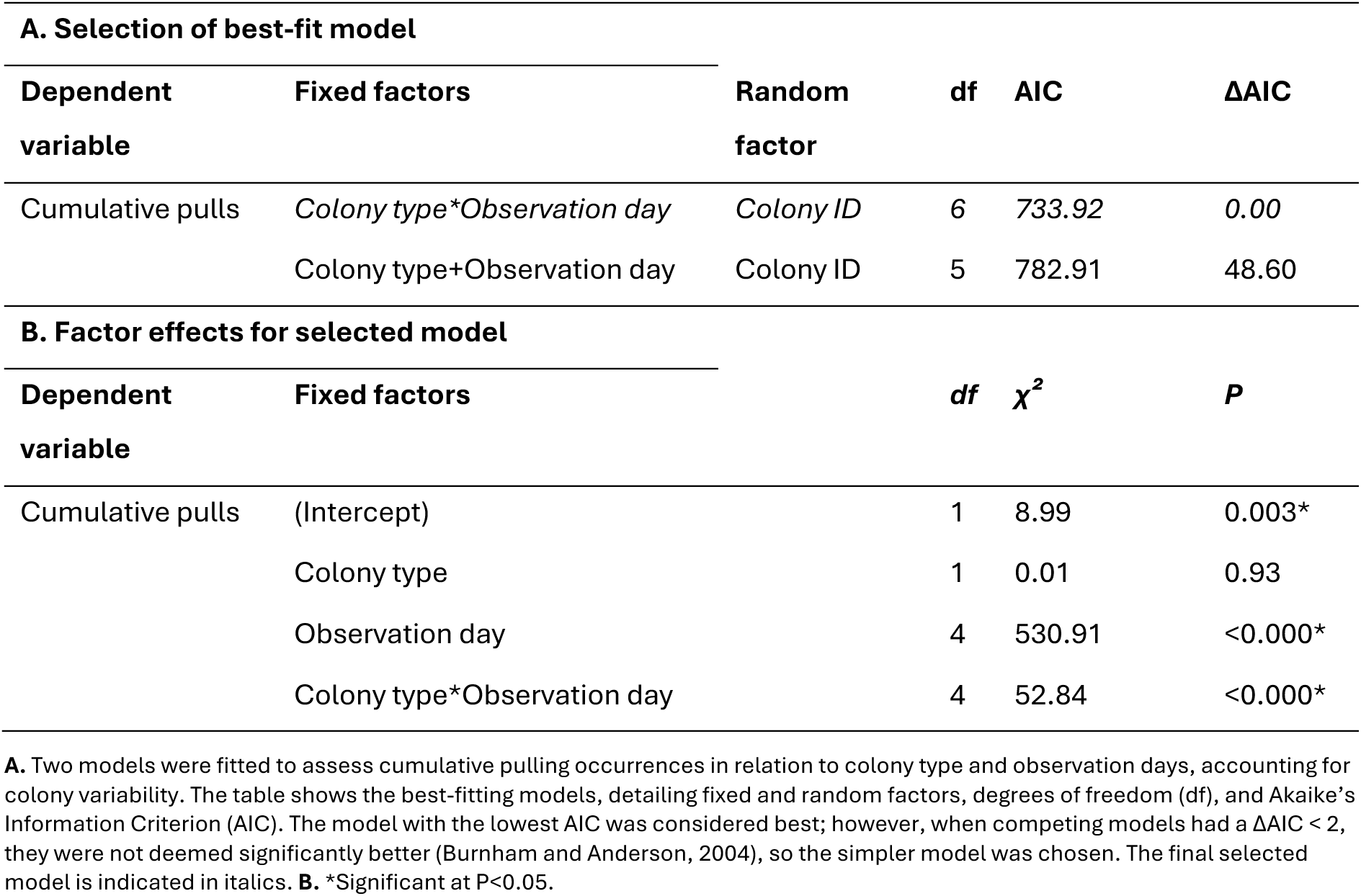
Comparison and summary of best-fit Poisson generalized linear mixed-effects models analyzing the impact of observation day and colony type (primary *Phase I* vs. secondary *Phase II*) on the cumulative string-pulling occurrences recorded.

| A. Selection of best-fit model |  |  |  |  |  |
| --- | --- | --- | --- | --- | --- |
| Dependent variable | Fixed factors | Random factor | df | AIC | $\Delta AIC$ |
| Cumulative pulls | *Colony type\*Observation day* | *Colony ID* | 6 | 733.92 | 0.00 |
|  | Colony type+Observation day | Colony ID | 5 | 782.91 | 48.60 |
| B. Factor effects for selected model |  |  |  |  |  |
| Dependent variable | Fixed factors | df | $\chi^2$ | P | |
| Cumulative pulls | (Intercept) | 1 | 8.99 | 0.003* |  |
|  | Colony type | 1 | 0.01 | 0.93 |  |
|  | Observation day | 4 | 530.91 | <0.000* |  |
|  | Colony type*Observation day | 4 | 52.84 | <0.000* |  |

A. Two models were fitted to assess cumulative pulling occurrences in relation to colony type and observation days, accounting for colony variability. The table shows the best-fitting models, detailing fixed and random factors, degrees of freedom (df), and Akaike's Information Criterion (AIC). The model with the lowest AIC was considered best; however, when competing models had a $\Delta AIC < 2$ , they were not deemed significantly better (Burnham and Anderson, 2004), so the simpler model was chosen. The final selected model is indicated in italics. B. \*Significant at $P < 0.05$ .

**Table S8.2.** Post hoc analysis of the interactions between colony type and observation day, and their impact on the cumulative string-pulling occurrences recorded.

| Observation day | Contrast | Estimate | SE | df | z | P |
| --- | --- | --- | --- | --- | --- | --- |
| 1 | Primary vs. Secondary | 0.09 | 0.94 | Inf | 0.10 | 0.925 |
| 2 | Primary vs. Secondary | 1.34 | 0.91 | Inf | 1.49 | 0.135 |
| 3 | Primary vs. Secondary | 1.79 | 0.90 | Inf | 1.98 | 0.048* |
| 4 | Primary vs. Secondary | 1.57 | 0.90 | Inf | 1.74 | 0.082 |
| 5 | Primary vs. Secondary | 1.70 | 0.90 | Inf | 1.88 | 0.060 |

\*Significant at $P < 0.05$ . The Bonferroni p-value adjustment method used was the R package emmeans. Primary colonies had had prior exposure to five days of open diffusion during *Phase I*, while secondary colonies were introduced at the start of *Phase II*.

**Table S9.1.**
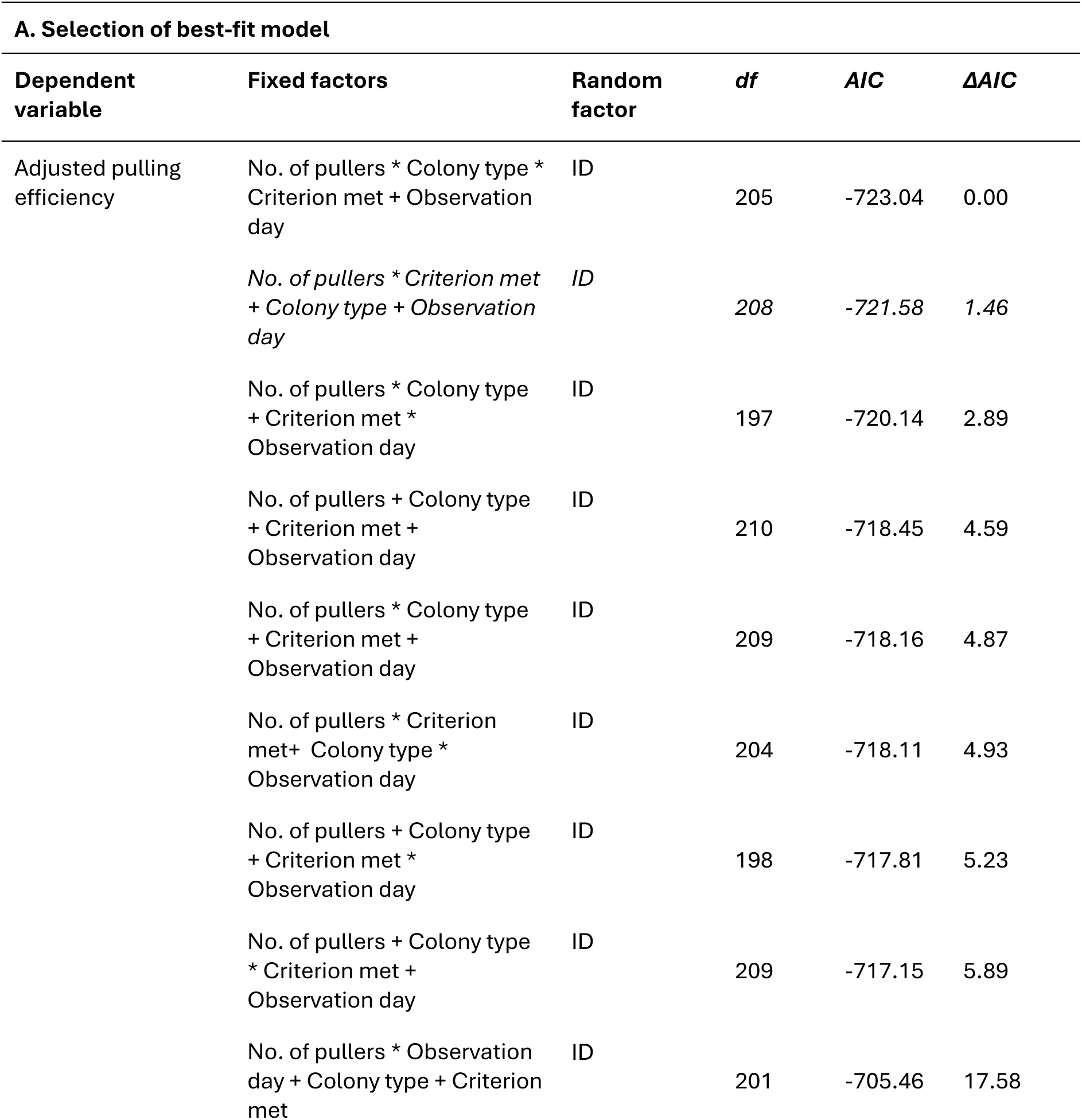
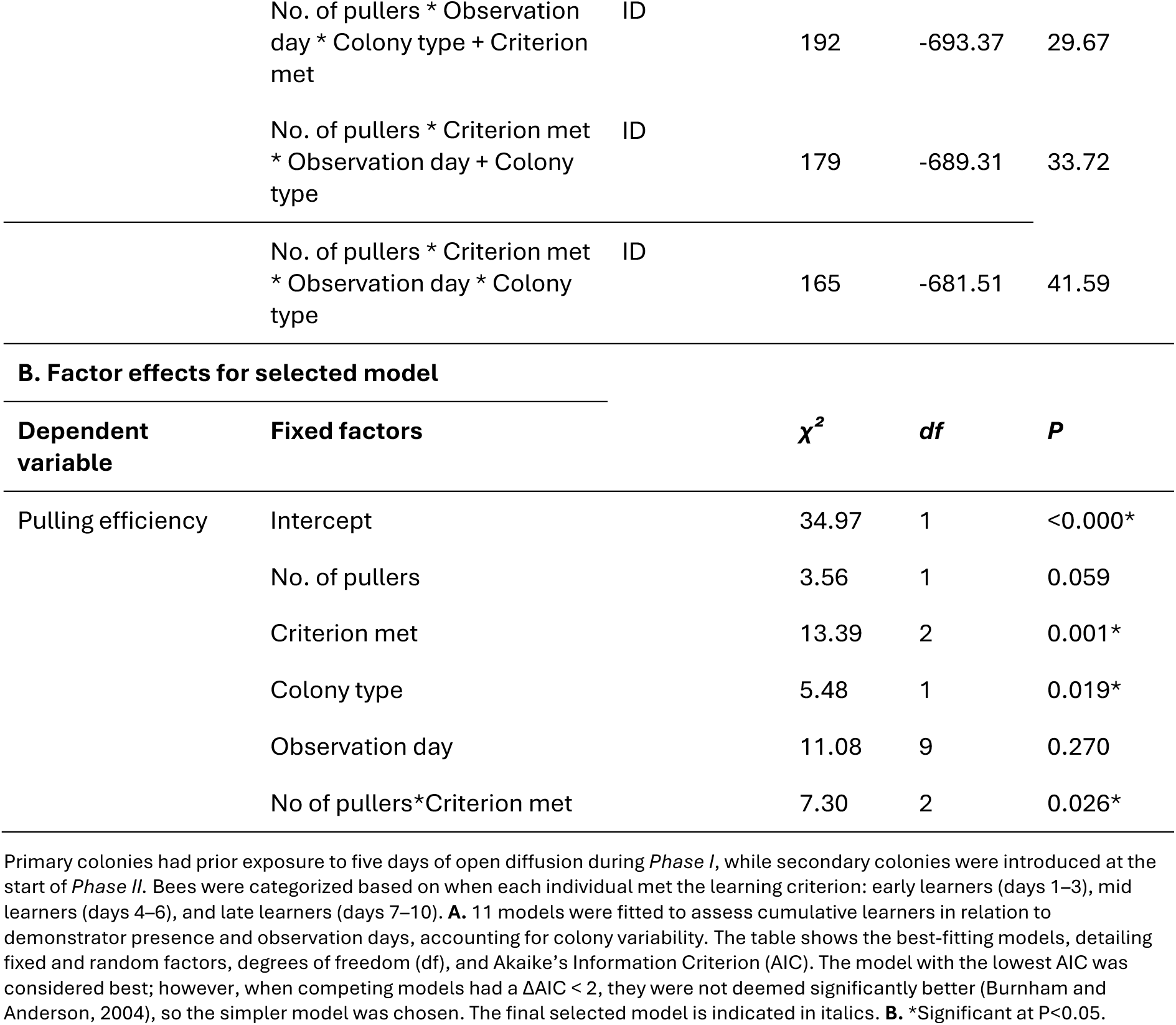
Summary of best-fit generalised mixed models analysing the impact of the number of bees recorded pulling the string per day (*no. of pullers*), colony type (*primary vs. secondary*), point of learning (*Criterion met: early, mid,* or *late*), and observation day on individual pulling efficiency across days, with control for individual variation.

**Table 9.2A.** Post hoc pairwise comparisons of estimated marginal means (EMMs) for the fixed effects listed in Table 7.1A.

| Significant fixed factor | Groups | Estimate | SE | df | z | P |
| --- | --- | --- | --- | --- | --- | --- |
| Colony type | Primary vs. Secondary | 0.69 | 0.30 | Inf | 2.34 | 0.019* |
| Criterion met | Early vs mid learners | 1.59 | 0.59 | Inf | 2.72 | 0.019* |
|  | Early vs late learners | 0.74 | 0.28 | Inf | 2.67 | 0.021* |
|  | Mid vs late learners | -0.85 | 0.60 | Inf | -1.41 | 0.335 |

\*Significant at $P < 0.05$ . The Bonferroni p-value adjustment method used was the R package emmeans. The table presents slope estimates (Estimate) for each group, along with standard errors (SE), z-scores (z), and p-values (P).

**Table 9.2B.** Post hoc analysis of the interaction between learner groups (Criterion met: early, mid, late) and the number of pullers (No. of pullers); see Table 7.1A.

| Group | Estimate | SE | Z | P |
| --- | --- | --- | --- | --- |
| Early | -0.07 | 0.04 | -1.89 | 0.059* |
| Mid | 0.03 | 0.05 | 0.68 | 0.496 |
| Late | 0.18 | 0.15 | 1.27 | 0.205 |

\*Significant at $P < 0.05$ . The Estimate column represents the slope of the relationship between the number of pullers and the proficiency (dependent variable) for each learner group (early, mid, late). Standard errors (SE), z-scores (z), and p-values indicate whether the slopes are significantly different from 0. The slopes were estimated using the R package emtrends.

**Table S10.**
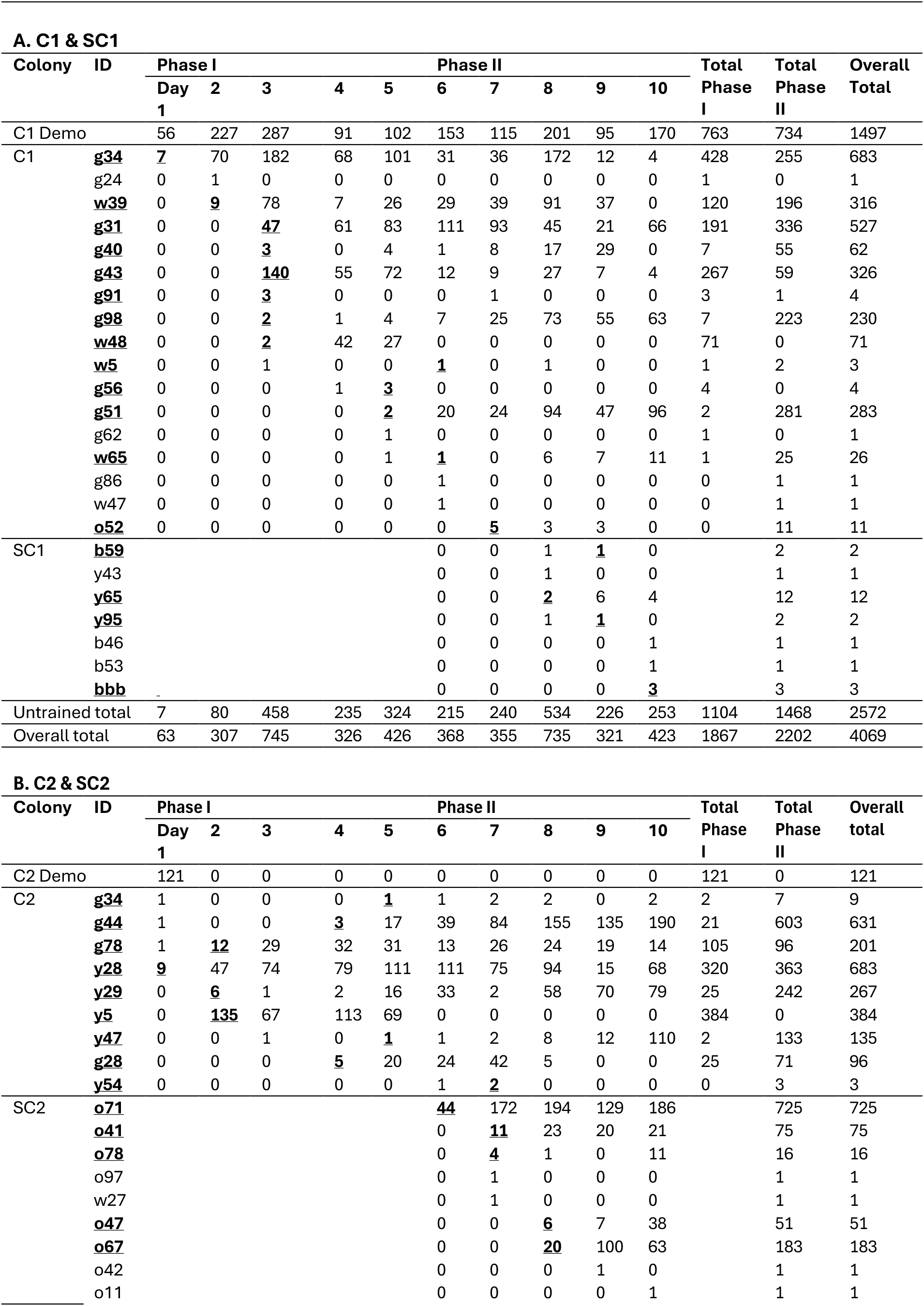
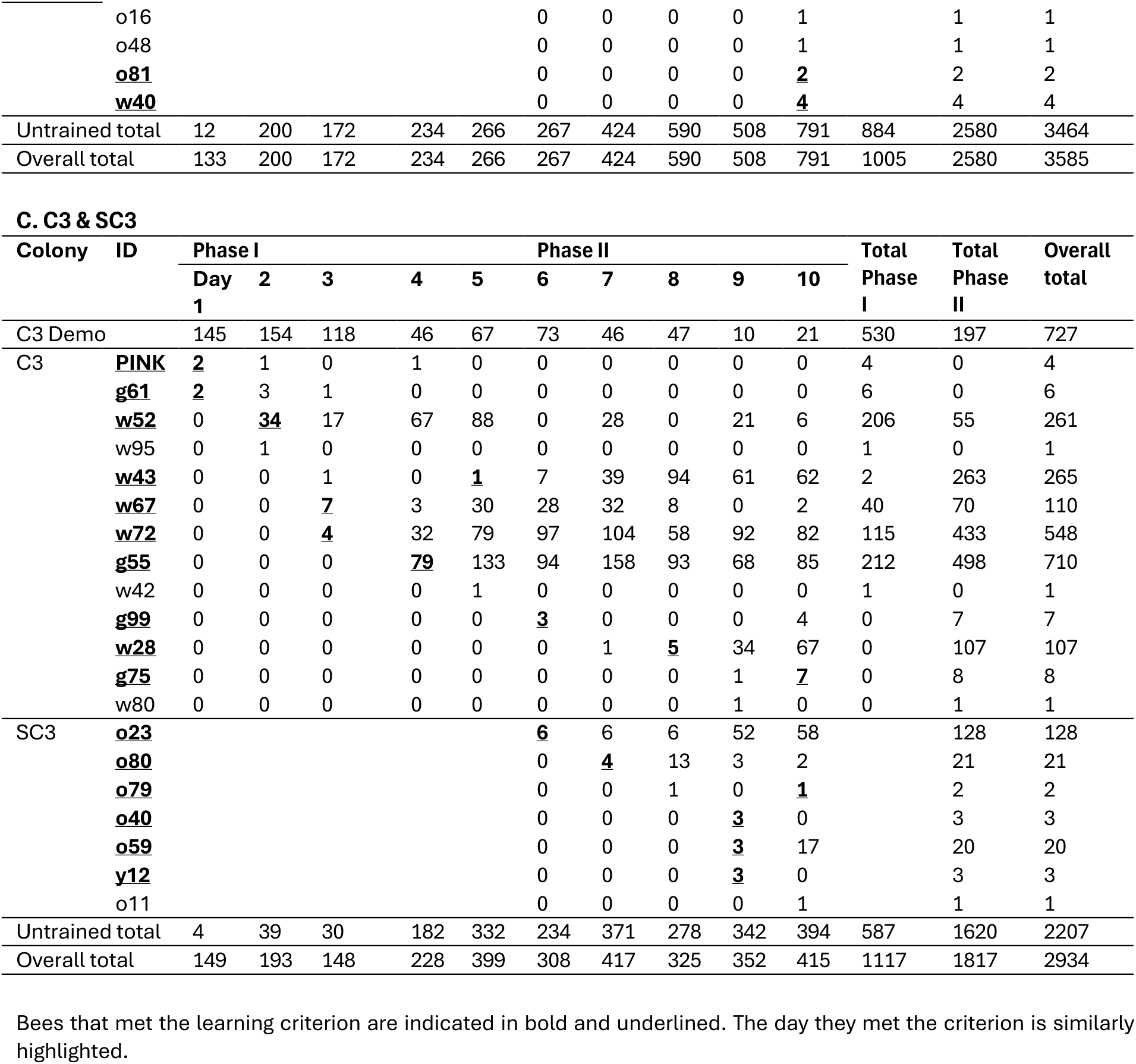
Daily string-pulling incidence for all bees in the experimental primary and secondary colonies.

**Table S11.** Daily string-pulling incidence for all bees in the control colonies.

| Colony | ID | Phase I |  |  |  |  | Total |
| --- | --- | --- | --- | --- | --- | --- | --- |
|  |  | Day 1 | 2 | 3 | 4 | 5 |  |
| CC3 | **g21** | 0 | 0 | 0 | <u>7</u> | 11 | 18 |
|  | g59 | 0 | 0 | 0 | 0 | 1 | 1 |
|  | **o94** | 0 | 0 | 0 | 0 | **4** | 4 |
| Untrained total |  | 0 | 0 | 0 | 8 | 17 | 25 |

Bees that met the learning criterion are indicated in bold and underlined. The day they met criterion is similarly highlighted.

**Supplementary Figure 1.**
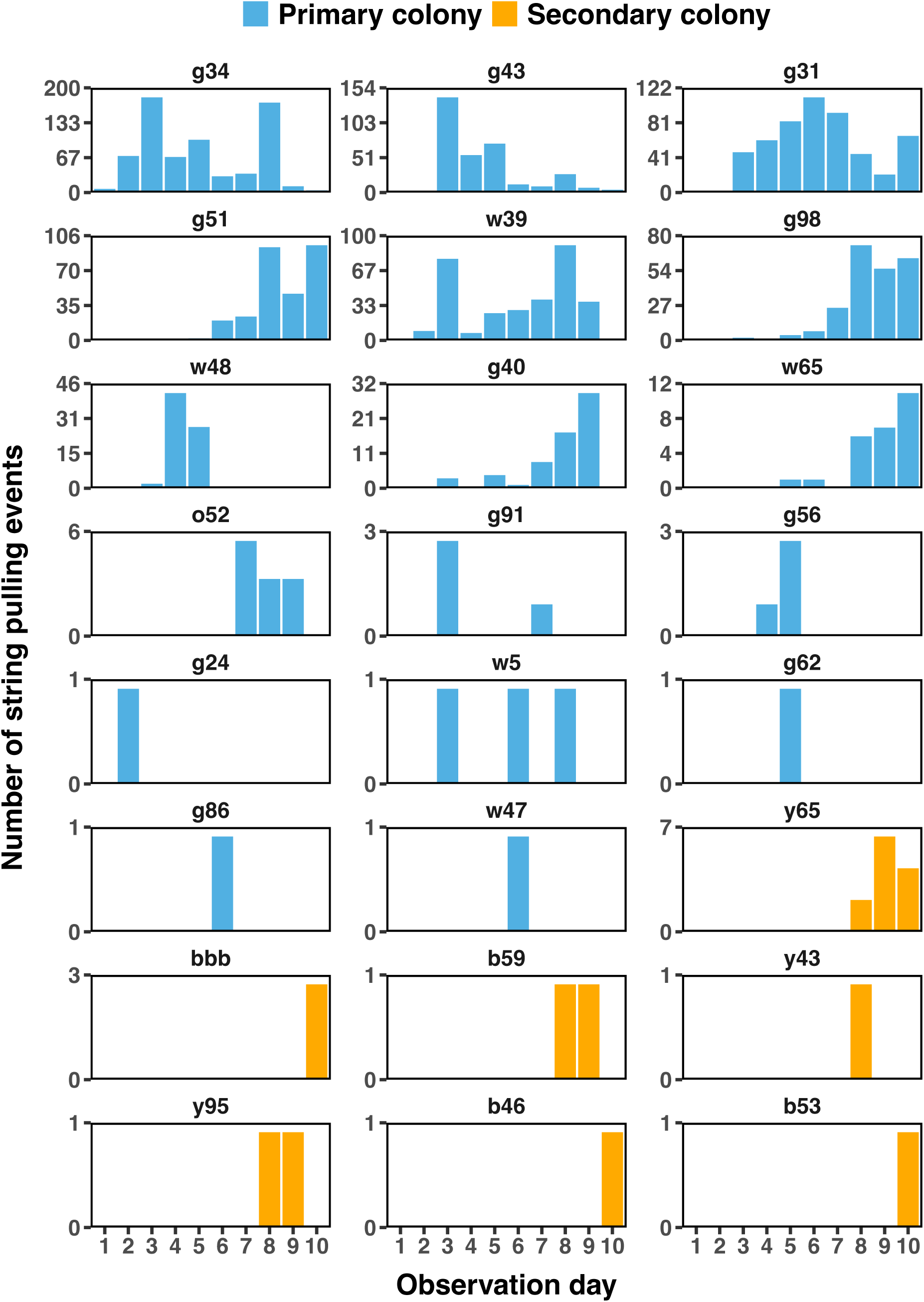
Daily string-pulling events for individual bees from the primary colony C1 (blue) and the secondary colony SC1 (yellow). Each bar represents the cumulative number of string-pulling events recorded for a specific bee across experimental days. The underlying raw data are presented in Supplementary Table S10.

**Supplementary Figure 2.**
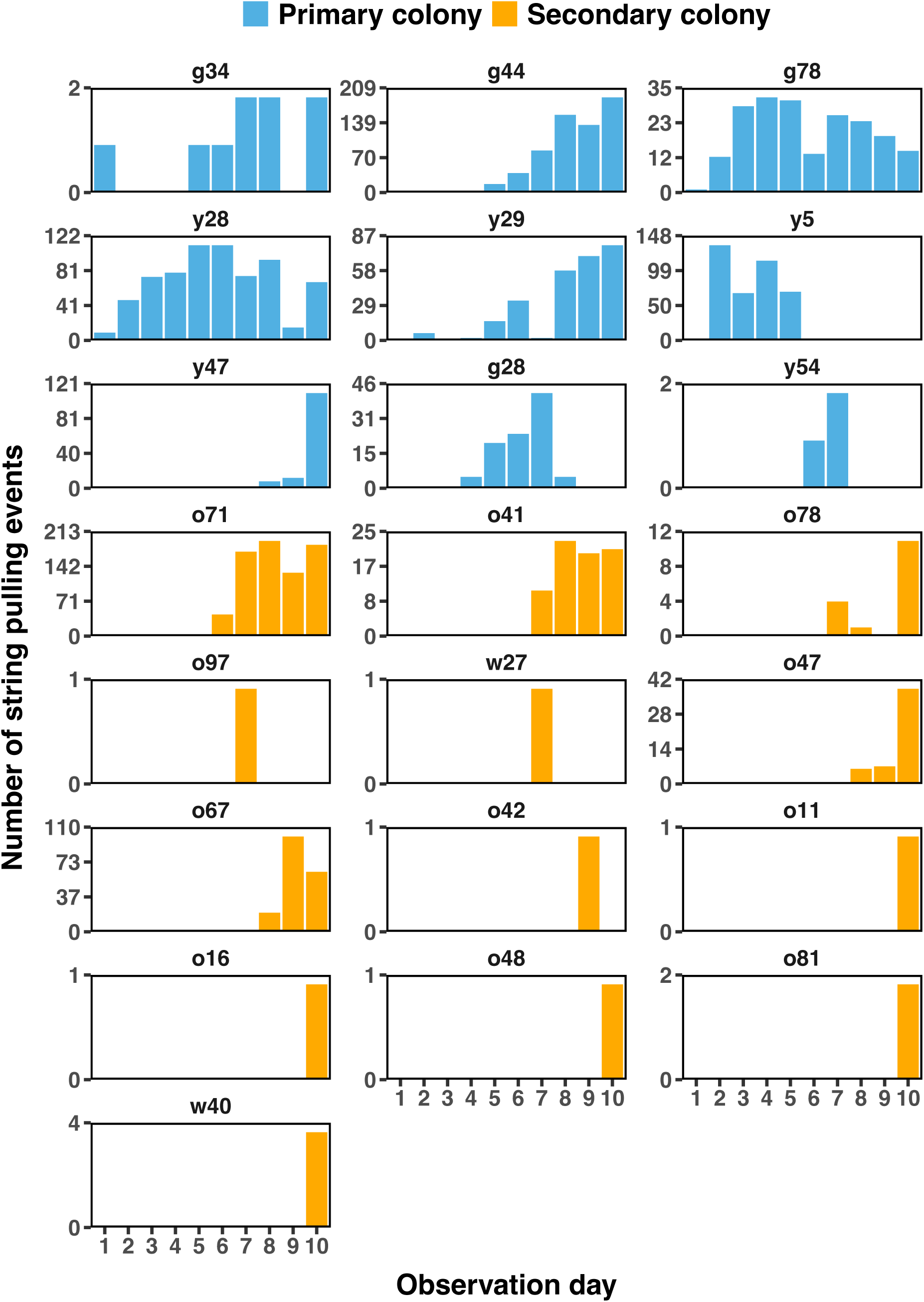
Daily string-pulling events for individual bees from the primary colony C2 (blue) and the secondary colony SC2 (yellow). Each bar represents the cumulative number of string-pulling events recorded for a specific bee across experimental days. The underlying raw data are presented in Supplementary Table S10.

**Supplementary Figure 3.**
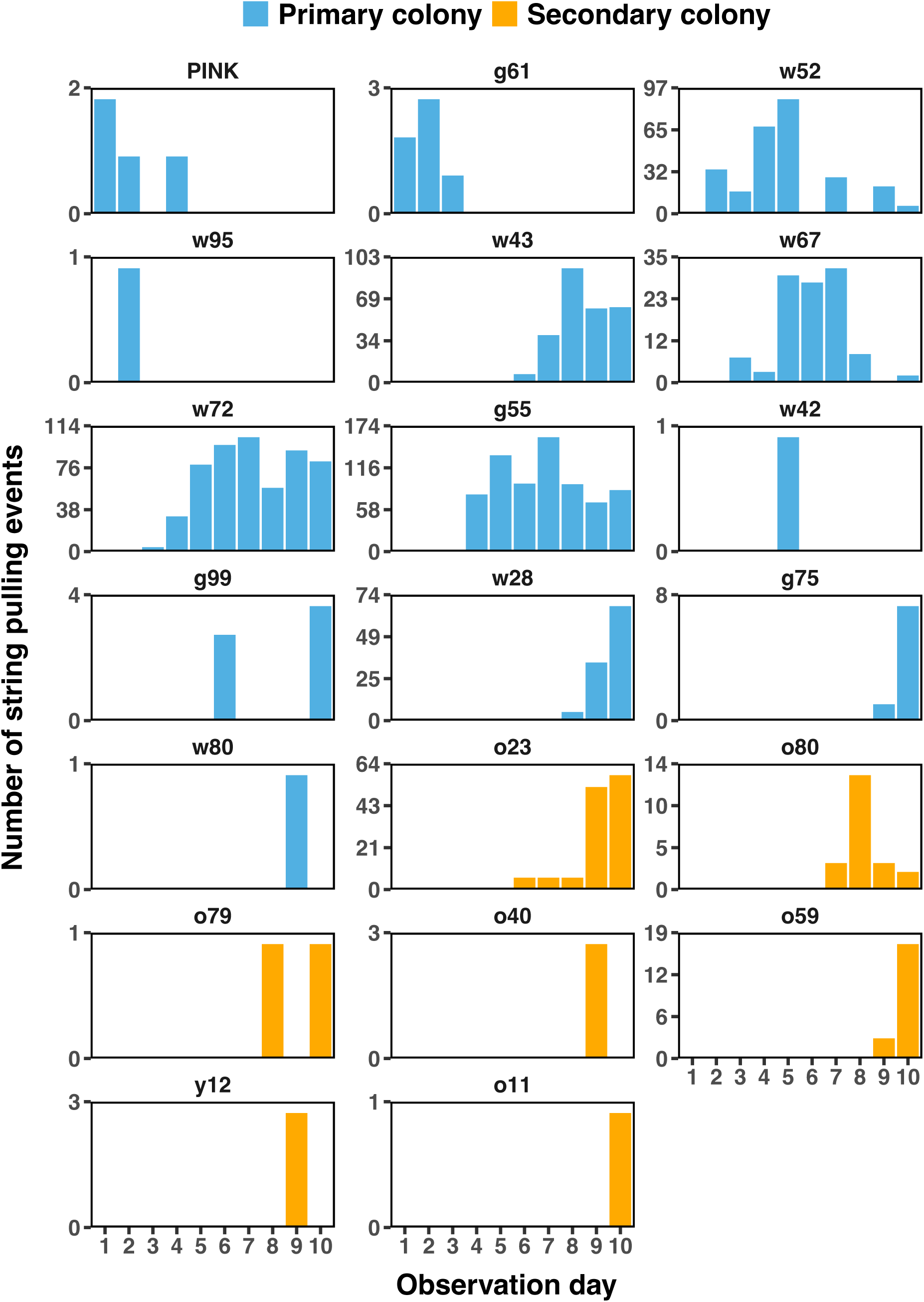
Daily string-pulling events for individual bees from the primary colony C3 (blue) and the secondary colony SC3 (yellow). Each bar represents the cumulative number of string-pulling events recorded for a specific bee across experimental days. The underlying raw data are presented in Supplementary Table S10.

**Supplementary Figure 4.**
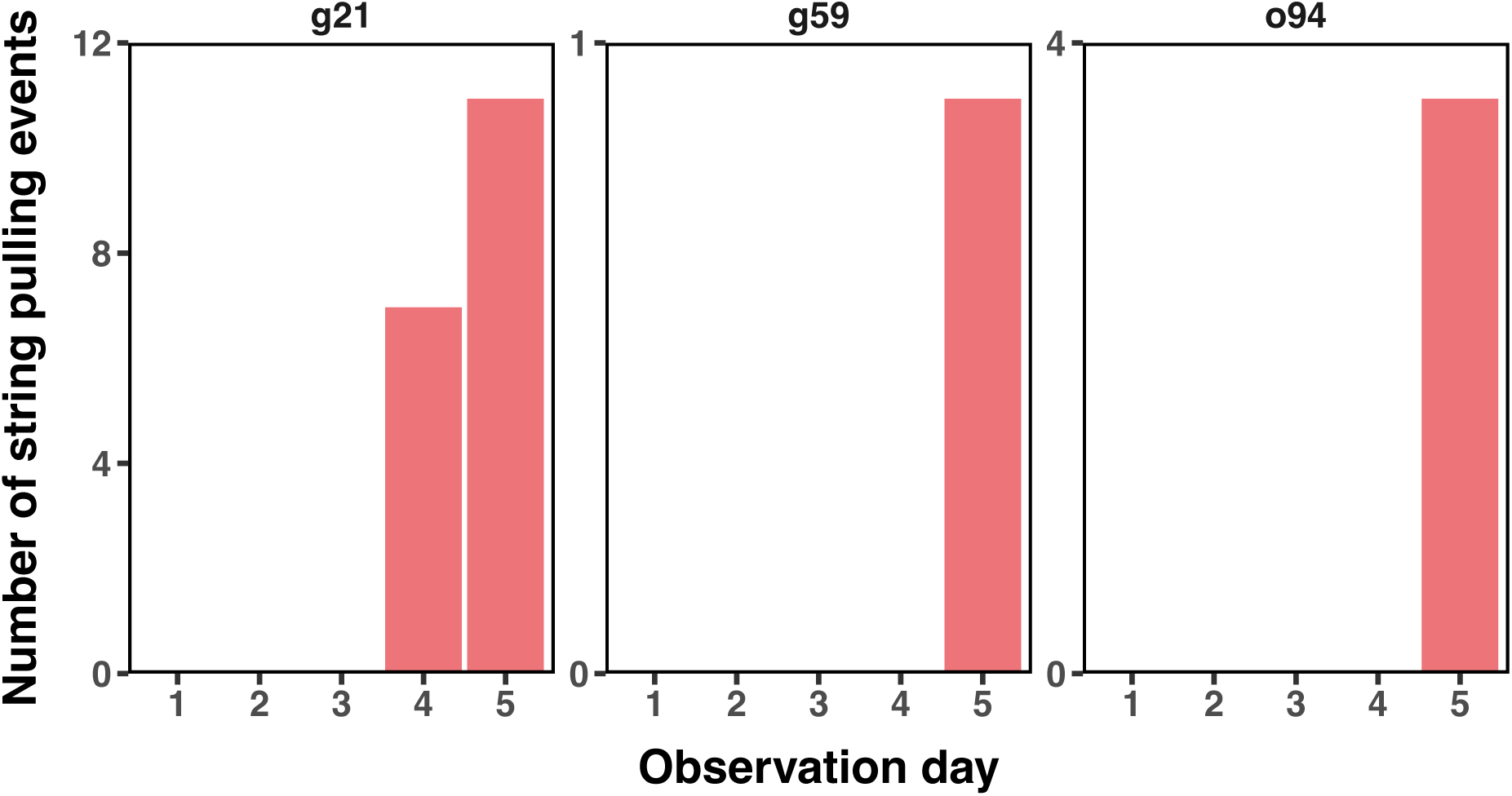
Daily string-pulling events for individual bees from the control colony CC3. Each bar represents the cumulative number of string-pulling events recorded for a specific bee across experimental days. The underlying raw data are presented in Supplementary Table S11.

**Supplementary Figure 5.**
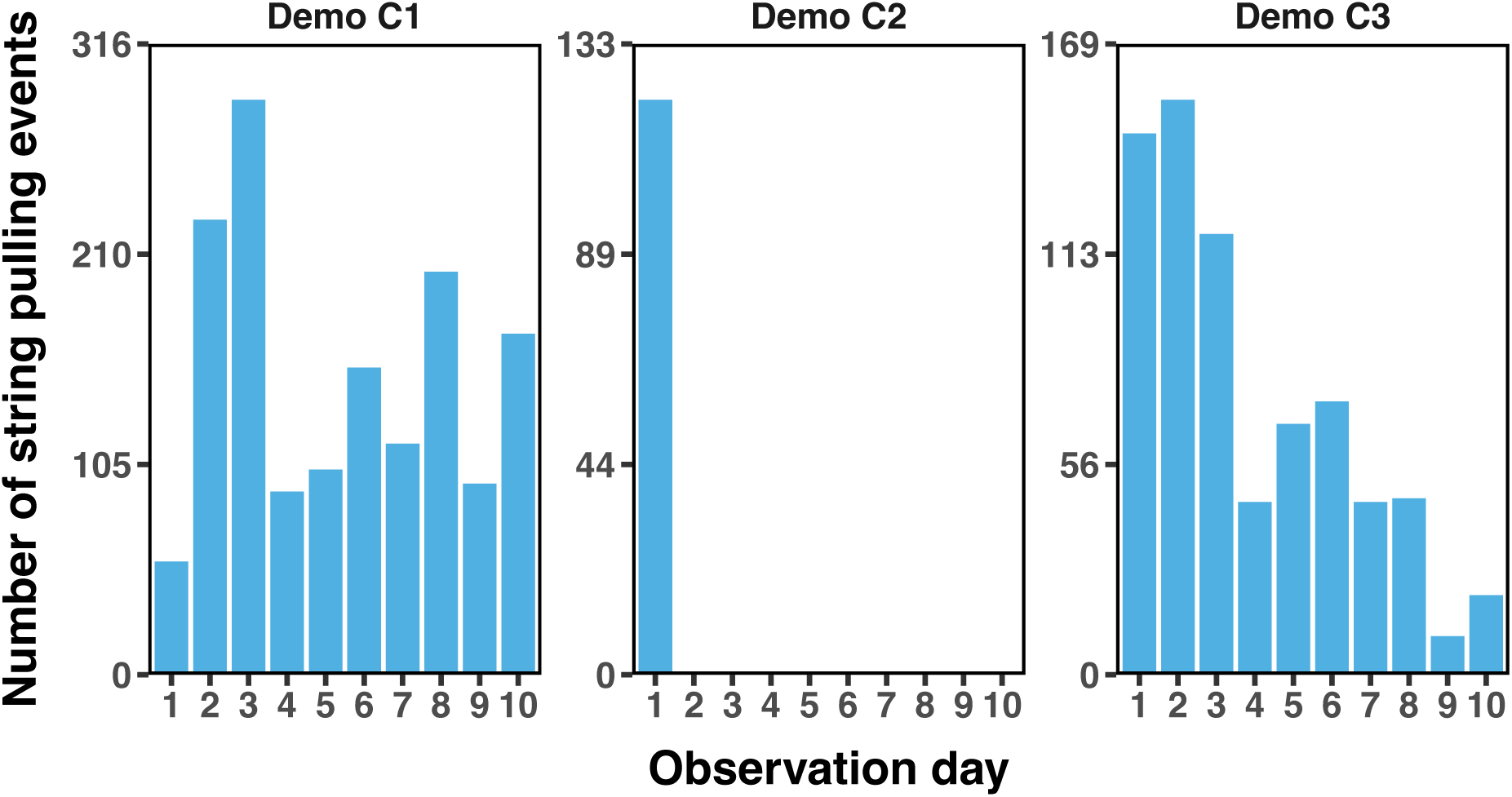
Daily string-pulling events for each seeded demonstrator. Each plot represents the number of string-pulling events performed by a single demonstrator bee across experimental days — experimental colony C1 (left), experimental colony C2 (middle), and C3 (right). The underlying raw data are presented in Supplementary Table S10.

**Supplementary Video 1. Footage of a sample string-pulling event performed by a learner.** Learner g31 (Experimental Colony 2) successfully extracts an artificial flower using the string-pulling technique to obtain a reward. A number of bees (in order of appearance: g40, w5, g58 and r44) attend to stages of the string-pulling behaviour. Once the flower is extracted, g40 scrounges the reward from g31.

**Supplementary Video 2. Footage of a sample string-pulling event performed by a learner**. Learner g55 (Experimental Colony 3) successfully extracts an artificial flower using the string-pulling technique to obtain a reward. A number of bees (in order of appearance: w28, g57, b12, w56 and w43) attend to stages of the string-pulling behaviour. Once the flower is extracted, w28 and w43 scrounge the reward from g55.

**Supplementary Video 3. Footage of a sample string-pulling event performed by a trained demonstrator.** Trained demonstrator y29 (Experimental Colony 1) successfully extracts an artificial flower using the string-pulling technique to obtain a reward. Another bee (w7) attends to part of the string-pulling behaviour.

**Supplementary Video 4. Footage of a flower extraction in the absence of string-pulling.** Multiple bees (g31, g86, w7, r52, w48, w39 and g34; Experimental Colony 1) contribute to the extraction of a single artificial flower and successfully obtain the reward, without showing any directed string-pulling behaviour. This and similar events were not included in the analysis.

## Notes

### Competing Interest Statement

The authors have declared no competing interest.

